# The Tip60 acetylome is a hallmark of the proliferative state in *Drosophila*

**DOI:** 10.1101/2025.07.15.664872

**Authors:** Zivkos Apostolou, Anuroop V. Venkatasubramani, Lara C. Kopp, Gizem Kars, Alessandro Scacchetti, Aline C. Sparr, Tamas Schauer, Axel Imhof, Peter B. Becker

## Abstract

The acetyltransferase KAT5/Tip60 is an epigenetic regulator of transcription and the DNA damage response. In *Drosophila*, Tip60 acetylates histones as part of the DOM-A complex, but it is unclear whether it has other substrates. In this work, we comprehensively studied the functions of Tip60 in a *Drosophila* proliferative cell model.

Depletion of Tip60 arrests the cell cycle, but remaining viable cells resist mutagenic irradiation. The impaired proliferation is explained by reduced expression of critical cell cycle genes. Tip60 binds their transcription start sites and Tip60-dependent acetylation of the histone variant H2A.V correlates with transcription activity.

A potentially synergistic pathway for cell cycle regulation involves the acetylation of proteins other than histones. The Tip60-dependent nuclear acetylome contains hundreds of proteins, many of which are involved in diverse aspects of cell growth and division, including replication, mitosis, gene expression, chromatin organization and ribosome biogenesis.

We hypothesize that Tip60 coordinates the proliferative state through histone and non-histone effectors. Reversible acetylation of diverse effector proteins bears potential for fine-tuning energy-intensive processes in response to stresses or nutritional shortcomings. Our study portrays the DOM-A/TIP60 complex as a general promoter of cell proliferation.

## Introduction

The acetylation of lysines in the N-terminal ‘tail’ domain of histones is an important principle of chromatin regulation (1). Lysine acetylation may disrupt the folding of the nucleosomal fiber or serve as binding site for bromodomain-containing epigenetic ‘reader’ proteins, which recruit additional coregulators (2–4).

The lysine acetyltransferase KAT5/Tip60 (Lysine Acetyltransferase 5/Tat-interacting protein of 60 kDa, (5)) is a powerful epigenetic regulator, best known for its ability to acetylate histones H4 and H2A. The importance of Tip60 is illustrated by the fact that its structure and essential functions in regulating transcription and the DNA damage response have been conserved during eukaryote evolution, albeit modified in interesting ways (6).

The orthologous acetyltransferase in *Saccharomyces cerevisiae*, Esa1, is the enzymatic subunit of the NuA4 complex (6–9). NuA4 activates transcription at least in part by stimulating the incorporation of the histone variant H2A.Z just downstream of transcription start sites (TSSs), a hallmark of active for promoters (10). The acetylation of H4 and H2A by NuA4 stimulates the SWR1 remodeling complex (SWR-c) to exchange H2A for the H2A.Z variant (11,12).

This functional interaction between NuA4 and SWR-c is structurally enforced in mammals, where the two paralogous complexes apparently fused to form a new entity, the p400 complex, combining KAT and nucleosome remodeling activities (13,14). This evolutionary merger presumably involved the fusion of the gene encoding the NuA4 scaffold subunit Eaf1 with a SWR1 paralog gene encoding a remodeling ATPase.

In *Drosophila*, most Tip60 resides in the Domino-A complex (15), a p400-like assembly that is scaffolded by the SWR1-related nucleosome remodeler, Dom-A. Tip60 is known to acetylate histone H4 at lysine 12 (H4K12ac) and lysines in the terminus of H2A.V (the H2A.Z variant in flies). Delivering Tip60 activity to promoters appears to be one major activity of the DOM-A complex, since so far, a histone exchange activity of DOM-A has not been detected (15).

In contrast to mammals, where the SWR1-type nucleosome remodeling enzymes p400 and SRCAP are encoded by different genes, the corresponding *Drosophila* proteins, Dom-A and Dom-B, arise as differential splice forms encoded by the *domino* gene (14,16).

Effects of Tip60 on cell cycle and the DNA damage response are likely to be mediated through gene regulation by histone acetylation at promoter-proximal nucleosomes (17). The mammalian TIP60 is a coactivator of transcription that cooperates with a range of prominent transcription factors to regulate cell cycle genes (18–20). Others concluded that TIP60 functions more as a global transcription activator found at most active gene promoters (21).

Here, we explore the role of Tip60 and its acetylation reactions in a well-established *Drosophila* proliferative cell model. Depletion of Tip60 or Dom-A leads to cell cycle arrest, which can be explained by reduced transcription of genes involved in proliferation control. The activity of these genes depends on binding of Tip60 binds to their promoters and acetylation of H2A.V in promoter-proximal nucleosomes. The acetylation of H4K12 is detected at putative enhancer elements.

While the activating roles of histones are well established, much less is known about the acetylation of non-histone substrates (22). We, therefore, explored the steady-state changes of the nuclear acetylome upon depleting Tip60 and identified hundreds of Tip60-dependent acetylation targets. The nature of these targets, combined with the gene expression analysis leads us to hypothesize a coordinating function of Tip60 in promoting cell growth, cell cycle progression and proliferation decisions.

## Materials and Methods

**Table 1.**
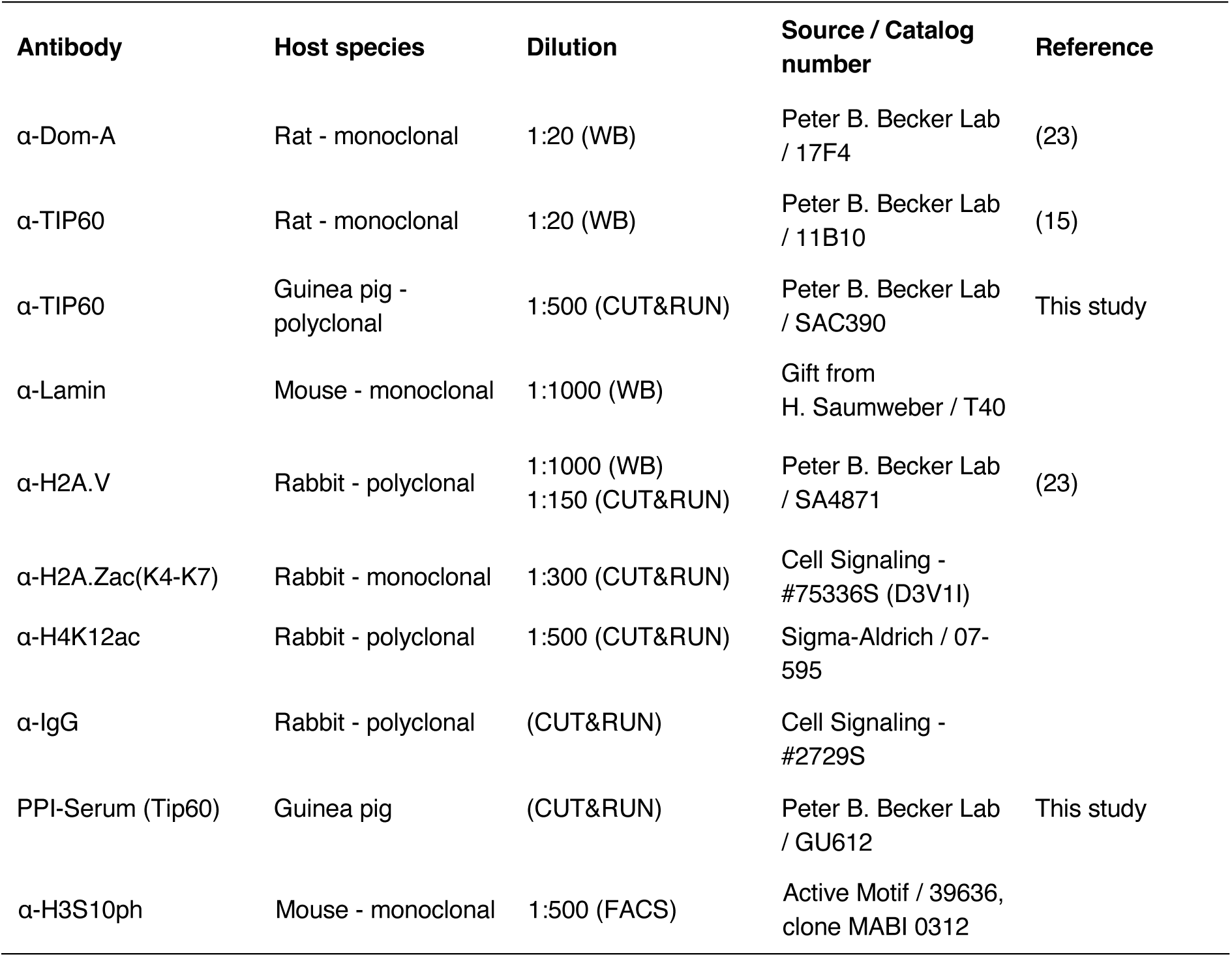
Antibodies.

### Cell lines and culture conditions

*Drosophila* embryonic Kc167 cells were obtained from *Drosophila* Genomic Resource Center. Cells were cultured at 26°C in Schneider’s *Drosophila* Medium (Thermo-Fischer, Cat. No. 21720024) supplemented with 10% FBS (Capricorn, Cat. No. FBS-12A) and 1% Penicillin-Streptomycin solution (PenStrep; Sigma-Aldrich, Cat. No. P-4333).

### RNA interference

Double-stranded RNA (dsRNA) targeting the desired sequences was generated by in vitro transcription using the HiScribe T7 High Yield RNA Synthesis Kit (NEB, Cat. No. E2040S). Primer sequences can be found in Supplementary Table 1.

For CUT&RUN, the cells underwent two consecutive rounds of RNAi. In brief, 6 μg of dsRNA was added to 10⁶ Kc167 cells in 0.5 ml of serum-free medium. Following a 1-hour incubation at 26°C, an equal volume of medium containing 20% fetal bovine serum (FBS) and 2% PenStrep was added, resulting in final concentrations of 10% FBS and 1% PenStrep. Cells were incubated in 12-well plates at 26°C for 3 days. On day 4, cells were split, counted and 0.8×10⁶ RNAi-treated cells of each condition was used for the second round of RNAi.

For the Proteome/Acetylome experiment, 180 μg of dsRNA was added to 3×10^7^ Kc167 cells in 15 ml of serum-free medium. Following a 1-hour incubation at 26°C, an equal volume of medium containing 20% fetal bovine serum (FBS) and 2% PenStrep was added, plus 20 ml of normal media to reach 50 ml of total volume, resulting in final concentrations of 10% FBS and 1% PenStrep. Cells were incubated in 175 cm^2^ cell culture flasks at 26°C for 6 days. In total, two 175 cm^2^ cell culture flasks were used per condition.

### Cell viability assay

Following three days of RNAi treatment, cells were counted, and 10⁶ cells were re-treated with dsRNA as previously described. The cells were then diluted 1:3 in fresh medium, and 50 μl (approximately 1.5×10⁴ cells) were seeded into white 96-well microplates with clear bottoms (BERTHOLD Technologies, Cat. No. 24910). On the following day (day 4), cells were exposed to varying doses of X-irradiation or UV-C irradiation and incubated at 26°C for an additional three days. Cell viability was assessed using the CellTiter-Glo Luminescent Cell Viability Assay (Promega, Cat. No. G7571), with luminescence signals measured on a Tecan Infinite M1000 microplate reader.

### Flow cytometry

#### RNAi interference

RNAi on Drosophila cells was performed using 6 μg of dsRNA, otherwise as described above. After three days of incubation, the cells were harvested and pelleted at 2000 rpm for 5 min. The number of cells in each sample was measured, and 10⁶ cells were re-suspended in 1 ml of serum-free Schneider’s *Drosophila* Medium supplemented with 1% PenStrep. The cells were re-seeded into 6-well plates and a second round of RNAi was performed using 10 μg of dsRNA identical to that used in the first round. The cells were then incubated for 1 hour and supplemented with 2 ml of complete medium. Following a further four days of incubation, the samples were prepared. Depending on the type of knockdown, cell confluency at the time of collection usually ranged between 60% and 85%.

#### Sample preparation

The cells were harvested and pelleted at 2000 rpm for 5 min., after which the pellet was washed twice with 1 ml of ice-cold PBS. After determining the cell count, 1.7×10⁶ cells from each sample were resuspended in 500 μl of ice-cold PBS in a 15 ml reaction tube. In non-sterile conditions, the cells were fixed by the dropwise addition of 4.5 ml of ice-cold 70% ethanol (Sigma-Aldrich, Cat. No. 32205-M), while continuously vortexing at a low speed. Samples were stored at −20°C for at least overnight and up to two weeks until analysis.

Unless stated otherwise, all subsequent steps were performed at 4°C with ice-cold reagents. All centrifugation and washing steps were performed at 2000 rpm.

The ethanol-fixed cells were pelleted for 15 minutes (min). After carefully decanting the supernatant, the cells were resuspended in PBS and left to rehydrate for 5 min. The cells were then washed twice with PBS for 5 minutes before being resuspended and permeabilized for 15 min in 1 ml of PBS containing 0.25% Triton X-100 (Sigma-Aldrich, Cat. No. T8787). The cells were pelleted, resuspended in 1 ml of PBS and distributed into 1.5 ml reaction vials. The cells were pelleted once and resuspended in 500 μl of 3% BSA in PBS at room temperature (RT). With the exception of the unstained and secondary antibody controls, the H3S10ph antibody (Active Motif, Cat. No. 39636, monoclonal antibody, mouse, clone MABI 0312) was added to the samples at a dilution of 1:500 and the samples were incubated on an overhead rotator for 2 hours at RT. The cells were then pelleted, washed once with PBS and resuspended in 500 μl of PBS/3% BSA. Except for the unstained and primary antibody controls, secondary anti-mouse antibody Alexa 555 was added to the samples at a dilution of 1:1000 and the samples were incubated on an overhead rotator for 1 hour at RT. From this point onwards, the samples were protected from light at all times.

Once more, the cells were pelleted, washed once with PBS and resuspended in 100 μl of PBS at RT. The DNA was then counterstained by adding 10 μl of 0.1 mg/ml DAPI (4’,6-diamidino-2-phenylindole, Sigma-Aldrich, Cat. No. No. 10236276001) directly to the cell suspension, except for the unstained and both antibody controls. After an incubation period of 10–15 min at RT, the samples were diluted by adding 100 μl of PBS and stored at 4°C until analysis. The samples were transferred into 5 ml Corning Falcon round-bottom tubes (Cat. No. 1172671, Omnilab-Laborzentrum GmbH) and the cell cycle profiles were measured using a low flow rate on a BD LSRFortessa Cell Analyzer (BD Biosciences). The 355 nm UV laser was used for DAPI (DNA content) and the 561 nm laser was used to visualize Alexa 555 bound to H3S10ph (mitotic index).

The results were analyzed using the FlowJoTM software (v10.8.1, BD Life Sciences).

### X-ray irradiation

The cells were irradiated with the indicated dose of grays in cell culture plates without lids, using a 130 kV, 5 mA Faxitron CellRad X-ray source.

### Cell fractionation

Cell fractionation was performed as previously described (24) with the following adjustments.

4×10^6^ of Kc167 cells were washed with PBS and resuspended in 200 μl of Buffer-A (10 mM HEPES pH 7.9 (KOH), 10 mM KCl, 1.5 mM MgCl_2_, 0.34 M sucrose, 10% glycerol, 1 mM DTT, 1x cOmplete EDTA-free Protease inhibitor (Roche, Cat. No. 5056489001), 1x PhosStop (Roche, Cat. No. 04906845001), 0.5 mM Na-butyrate) with the addition of 0.1% Triton-X-100, and incubated at 4°C for 10 min. From here on all procedures at 4°C. Nuclei were pelleted at 1500 g for 4 min (pellet: P1). The supernatant (S1) was centrifuged at 13000 g for 10 min to remove cell debris and insoluble aggregates (S2). The nuclear pellet P1 was washed with Buffer-A and resuspended nuclei in 200 μl of Buffer-B (3 mM EDTA, 0.2 mM EGTA, 1 mM DTT, Protease inhibitor, 1x PhosStop, 0.5 mM Na-butyrate) for lysis and incubated for 10 min. Samples were centrifuged at 2000 g for 4 min to separate soluble nuclear proteins (S3) from chromatin (P3). P3 was washed with Buffer-B and spun down at 13000g for 1 min. The chromatin pellet (P3) was resuspended in an equal volume equivalent to S3 in laemmli sample buffer (LSB; 50 Tris pH 6.8, 10% glycerol, 2% SDS, 0.01% bromophenol, 0.1M DTT) and boiled for 10 min.

### Antibodies

A polyclonal antibody against Tip60 was generated by expressing the full-length Tip60 protein as a N-terminal glutathione-S-transferase (GST) fusion in *E. coli*. The recombinant protein was purified using glutathione sepharose resin (GE Healthcare, Cat. No. 17075605) and subsequently eluted with glutathione. Antibody production in guinea pigs was outsourced to Eurogentec (https://secure.eurogentec.com/eu-home.html). The Tip60 antibody was validated through RNA interference (RNAi) experiments followed by western blot analysis.

### Proteome and acetylome

#### Extraction of nuclear proteins

Cells of 2 big flasks 175 cm^2^ were collected in one Corning bottle and pelleted at 1000 g for 15 min. Cells were resuspended in 10 ml of PBS and pelleted in 15 ml falcons at 1000 g for 5 min. This was followed by resuspension of pellet in 10 ml NBT-10 buffer (10% sucrose, 0.15% Triton, 0.5 mM EGTA pH 8, 60 mM KCl, 15 mM NaCl, 15 mM HEPES pH 7.5, 1 mM PMSF, 1 mM DTT, 1X cOmplete EDTA-free Protease Inhibitor-Roche, 1 mM Na-butyrate) and rotated on a wheel for 10 min at 4°C. The content of each tube was added on top of 20 ml cushion NB-1.2 buffer (1.2 M sucrose, 0.5 mM EGTA pH 8, 60 mM KCl, 15 mM NaCl, 15 mM HEPES pH 7.5, 1 mM PMSF, 1 mM DTT, 1X cOmplete EDTA-free Protease Inhibitor-Roche, 1 mM Na-butyrate) and the nuclei was pelleted by centrifuging at 4000 rpm for 20 min. The nuclear pellet was resuspended in 1 ml of NBT-10 buffer and put in 15 ml falcon followed by another centrifugation at 4000 rpm for 10. This was followed by a brief wash with cold PBS and another centrifugation at 4000 rpm for 5. Nuclei pellets were resuspended in 2 ml Urea Lysis Buffer (ULB: 20 mM HEPES pH 8.0, 9 M urea, 1 mM sodium orthovanadate, 2.5 mM sodium pyrophosphate, 1 mM β-glycerophosphate, 1 mM Na-butyrate, 60 μM Sirtinol), incubated on ice for 10 min and snap-frozen in liquid-N2.

#### Acetylome sample preparation

Sample preparation for acetylome was performed with PTMScan® Acetyl-Lysine Motif [Ac-K] kit (Cell Signaling, 13416) and manufacturer’s protocol was followed. To the obtained nuclear extract, 9 ml of ULB was added and sonicated in ice with a tip sonifier (Branson, Model-250D) at 40% amplitude with 15 sec ON and 50 sec OFF for 3-5 times. To the sonified lysate, 1/278^th^ volume of 1.25 M DTT was added and incubated at RT for 60 min, followed by 15 min incubation in dark with 1/10^th^ volume of iodoacetamide (Merck, 8.04744.0025). The supernatant was then diluted with 20 mM HEPES pH 8.0 and incubated (with mixing) with 1 mg/ml trypsin (Sigma-Aldrich, 175641-5G) at RT for overnight. Trypsin was added at 1:100 ratio with total initial protein amount. Following overnight incubation, the digestion was confirmed with SDS-PAGE and 1/20 volume of 20% TFA was added to the digested peptide solution and incubated for 15 min on ice. The lysate was then centrifuged at 1,780 g for 15 min at RT to remove any precipitate.

For peptide purification, Sep-Pak^®^ Light C18 cartridges filter column (Waters, WAT023501) was connected to a 10cc syringe and the column was pre-wetted with 5 ml 100% ACN, followed by sequential washes with 1 ml, 3 ml and 6 ml of 0.1% TFA. Acidified digest was then loaded on to the column (without vacuum), followed by further washes of 1 ml, 5 ml and 6 ml with 0.1% TFA and then with 2 ml of wash buffer (0.1% TFA and 5% ACN). Elution of the peptide was carried out by washing the column with 0.1% TFA and 40% ACN. Eluted peptide was then frozen overnight at −80°C and lyophilized for at least 48 hours.

Lyophilized peptide was centrifuged at 2,000 g for 5 min at RT and resuspended with 1.4 ml of the IAP buffer (provided by the manufacturer) and centrifuged again at 10,000 g at 4°C for 5 min. 10% of the cleared solution was taken as input for proteome. Lysine antibody beads (provided by the manufacturer) were washed with 1 ml of PBS, centrifuged at 2,000 g, resuspended with 40 μl of PBS. To the cleared solution, half the recommended volume of antibody-bead slurry was used and incubated in a rotator for 2 hr at 4°C. Incubated samples were then centrifuged at 2,000 g for 30 s, supernatant transferred to a new vial for further use. Beads were then washed twice with 1 ml of IAP buffer, mixed by inverting, centrifuged at 2,000 g for 30 sec, followed by similar washes, thrice with 1 ml chilled HPLC water. Peptides bound to the beads were eluted with twice by adding 55 μl of 0.15% TFA, vortexing, incubating for 10 min at RT and centrifuging for 30 sec at 2,000 g.

Desalting of the eluted peptide was performed with AttractSPE^®^ Disks Tips C18 column (affinisep, Tips-C18.T2.200.96). The column was first equilibrated with 50 μl 0.1% TFA twice. Input proteome and IP sample was then added to the C18, followed by washes with 0.1% TFA twice. Peptides were eluted with 10 μl of 0.1% TFA and 40% ACN, dried with vacuum concentrator and resuspended with 15 μl of 0.1% TFA for mass spectrometry analysis.

#### Mass spectrometry

Samples were evaporated to dryness, resuspended in 15 μl of 0.1% formic acid solution and injected in an Ultimate 3000 RSLCnano system (Thermo) separated in a 25-cm Aurora column (Ionopticks) with a 100-min gradient from 6-43% of 80% acetonitrile in 0.1% formic acid. The effluent from the HPLC was directly electrosprayed into a Orbitrap Exactive 480 (Thermo) operated in data dependent mode to automatically switch between full scan MS and MS/MS acquisition. Survey full scan MS spectra (from m/z 350-1200) were acquired with resolution R=60,000 at m/z 400 (AGC target of 3×10^6^). The 20 most intense peptide ions with charge states between 2 and 6 were sequentially isolated to a target value of 1×10^5^, and fragmented at 30% normalized collision energy. Typical mass spectrometric conditions were: spray voltage, 1.5 kV; no sheath and auxiliary gas flow; heated capillary temperature, 275°C; intensity selection threshold, 3×10^5^.

## CUT&RUN

CUT&RUN was performed as previously described (25). Briefly, Concanavalin-A magnetic beads (10 μl of slurry/reaction) were pre-activated by washing 2 times with Binding Buffer (BB: 20 mM Hepes pH 7.5 (NaOH), 10 mM KCl, 1 mM CaCl_2_, 1 mM MnCl_2_). 10^6^ Kc167 cells per condition were bound to the beads, and washed twice with Wash Buffer (WB: 20 mM Hepes pH 7.5 (NaOH), 150 mM NaCl, 0.5 mM spermidine, 1x cOmplete EDTA-free Protease inhibitor). Cells were permeabilized using 150 μl of Antibody Buffer (AB: 20 mM Hepes, pH 7.5 (NaOH), 150 mM NaCl, 0.5 mM spermidine, 0.05% digitonin, 2 mM EDTA, 0.5 mM Na-butyrate). Antibodies were mixed into AB at the indicated dilutions. Samples were put on a nutator at RT for 2 hours, and washed twice with Dig-Wash Buffer (Dig-WB: 20 mM Hepes, pH 7.5 (NaOH), 150 mM NaCl, 0.5 mM spermidine, 0.05% digitonin, 1x cOmplete EDTA-free Protease inhibitor). pAG/MNase protein (700 ng/ml) was mixed into 150 μl of Dig-WB and samples were placed on a nutator at 4°C for 1 hour, and then washed twice with Dig-WB. Chromatin digestion was performed by the addition of 2 mM (f.c.) CaCl_2_ into 100 μl of Dig-WB and incubated at 4°C for 30 min. The reaction was stopped by adding 100 μl of 2x STOP buffer (340 mM NaCl, 20 mM EDTA, 4 mM EGTA, 0.05% digitonin, 100 μg/ml RNase A, 50 μg/ml glycogen) and the chromatin fragments were released from insoluble nuclear chromatin by incubating at 37°C for 30 min. Samples were collected by a quick spin and placed on a magnet stand to concentrate the magnetic beads with bound chromatin fragments. Chromatin was digested with 0.25 mg/ml Proteinase K/ 0.1% SDS, and DNA was purified by phenol/chloroform extraction. DNA concentrations were determined using Qubit (Thermo Fisher).

### Library preparation and sequencing

For the CUT&RUN samples, sequencing libraries were prepared using NEB Next Ultra II DNA Library (New England Biolabs) as described (25). All libraries were sequenced on an Illumina NextSeq1000/2000 sequencer at the Laboratory of Functional Genomic Analysis (LAFUGA, Gene Center Munich, LMU). For CUT&RUN about 5×10^6^ paired-end reads were sequenced per sample for each of the biological replicates.

### Data analysis

#### Transcriptome

Analysis of raw reads for RNA sequencing data, PCA and scatter plots was adopted from Scacchetti et al. (15). Scripts are available on GitHub (https://github.com/tschauer/Domino_RNAseq_2020). Over-representation analysis, was performed using the org.Dm.eg.db (v3.21.0) (26) database and the enrichGO function from clusterProfiler (v4.16.0) (27). Genome alignment and gene and transcript quantification for Figure 5E were performed using the *Drosophila melanogaster* genome assembly BDGP6.46 (Ensembl), with STAR (v2.7.1a) (28) for alignment and RSEM (v1.3.0) (29) for quantification. All plots were generated using R graphics (30).

#### Proteome and acetylome data analysis

Raw mass spectrometry data files were processed using MaxQuant (31) (v2.1.3.0) with the *Drosophila melanogaster* reference proteome fasta obtained from UniProt. The resulting output files — “*proteinGroups.txt”* for proteome data and “*Acetyl(K)Sites.txt”* for acetylome data — were analyzed in the R environment (30), using the RAmP package (v1.3.5) available at (https://github.com/anuroopv/RAmP). Differential expression analysis (DEA) was performed using default parameters, with the following modifications: for the proteome dataset, RAmP::DEA(prot.Data = proteinGroups, sampleTable = SampleTable, org = “dme”, quantification = “LFQ”, contrast = c(”tip60_0_vs_GST_0”, “chm_0_vs_GST_0”), Fraction = “Proteome”, filter.protein.type = “fraction”, filter.protein.min = 0.75, impute.function = “bpca”, enrich = “ora”, ont = “BP”, simplify = TRUE, simplify_cutoff = 1, plotType = “treePlot”); for the acetylome dataset, RAmP::DEA(prot.Data = proteinGroups, enrich.Data = Acetyl(K)sites, sampleTable = SampleTable, org = “dme”, quantification = “LFQ”, contrast = c(”tip60_0_vs_GST_0”, “chm_0_vs_GST_0”), Fraction = “Enriched”, filter.protein.type = “fraction”, normalize = TRUE, filter.protein.min = 0.75, impute.function = “bpca”, enrich = “ora”, ont = “BP”, simplify = TRUE, simplify_cutoff = 1, plotType = “treePlot”)

For both the proteome and acetylome datasets, proteins detected in at least 75% of all conditions were initially filtered and retained for downstream analysis. These were classified as ‘Class 2’. After filtering, missing values were imputed using the BPCA algorithm, followed by differential expression analysis performed with the limma package (32). For each replicate, acetylome intensities were normalized to the mean LFQ intensity of the corresponding proteome condition. Significance was determined using both q-values and Π-values (33,34), with a cut-off of 0.05 applied in both cases. Proteins that failed to meet the 75% detection threshold in at least one condition were classified as ‘Class 1’. A protein was assigned to ‘Class 1’ if it exhibited 2-4 missing values (NAs) in one experimental condition while having 0-2 NAs in the other, based on the following allowed combinations: 4-0, 4-1, 4-2, 3-0, 3-1, and 2-0.

For the over-representation analysis, the entire quantified proteome served as the background for proteome data, while the full set of identified acetylated proteins was used as the background for acetylome analysis. Significant proteins from the proteome and acetylome analyses were separated by log-fold change to distinguish upregulated and downregulated (or differentially acetylated) peptides and their corresponding proteins. Gene Ontology (GO) term enrichment was then performed separately for each group, applying an FDR < 0.05 threshold for significance. Both ‘Class 1’ and ‘Class 2’ proteins were included in the over-representation analysis.

### CUT&RUN

Raw reads from FASTQ files were first trimmed to remove sequencing adaptors using cutadapt (v1.16) (35). Trimmed reads were then aligned to the *Drosophila melanogaster* reference genome (Ensembl BDGP6.46) using Bowtie2 (36,37), followed by processing with SAMtools (v1.9) (38) to filter alignments with a mapping quality of at least 2 (-q 2), and BEDTools (v2.28.0) (39). BigWig files were generated using bamCompare (deepTools v3.5.0) (40), normalizing each CUT&RUN sample to its corresponding IgG control (for H2A.V and H2A.Vac) or pre-immune serum (for Tip60), while also adjusting for sequencing depth. For each condition, biological replicates were averaged according to the number of mapped reads using bigwigAverage (deepTools v3.5.6) (40). All fragment sizes were used for further analysis. Peak calling for Tip60 averaged replicates was performed with MACS3 (41) using the parameters: cutoff 5, min-length 100, and max-gap 500. Custom code for CUT&RUN analysis is available on Zenodo (https://doi.org/10.5281/zenodo.15844155).

### Plots and statistical analysis

All plots, graphs and statistical analyses were performed in the R environment (30) unless stated otherwise. The choice of statistical tests was based on the experimental design and data type and is indicated in the corresponding figure legends.

## Results

### The DOM-A/TIP60 complex is necessary for cell proliferation and cell cycle progression

To investigate the role of the DOM-A/TIP60 complex in cell proliferation and cell cycle progression in *Drosophila* Kc167 cells, we reduced the expression of *tip60*, *dom-A* or *dom-B* through RNA interference (RNAi) alongside an RNAi control directed at irrelevant sequences of glutathione-S-transferase (*gst*) of *Schistosoma japonicum*. The knockdown efficiency was confirmed by immunoblot analysis (Supplementary Figure S1A), demonstrating efficient depletion of both Dom-A and Tip60 proteins. As observed before (15), complex disruption by depletion of Dom-A led to reduction of Tip60, but not vice versa.

Cell viability was assessed using a luminescence-based CellTiter-Glo assay on day 4 and day 7 of RNAi treatment. As shown in Figure 1A, depletion of Dom-A and Tip60 led to markedly reduced viable cell numbers. *Dom-A* knockdown resulted in more pronounced effects, indicating that *Dom-A* may have additional functions beyond serving as a delivery platform for Tip60. By comparison, depletion of Dom-B, the main H2A.V exchange factor (15), also impaired cell viability, though to a lesser extent. Control cells proliferated similarly to untreated cells.

**Figure 1.**
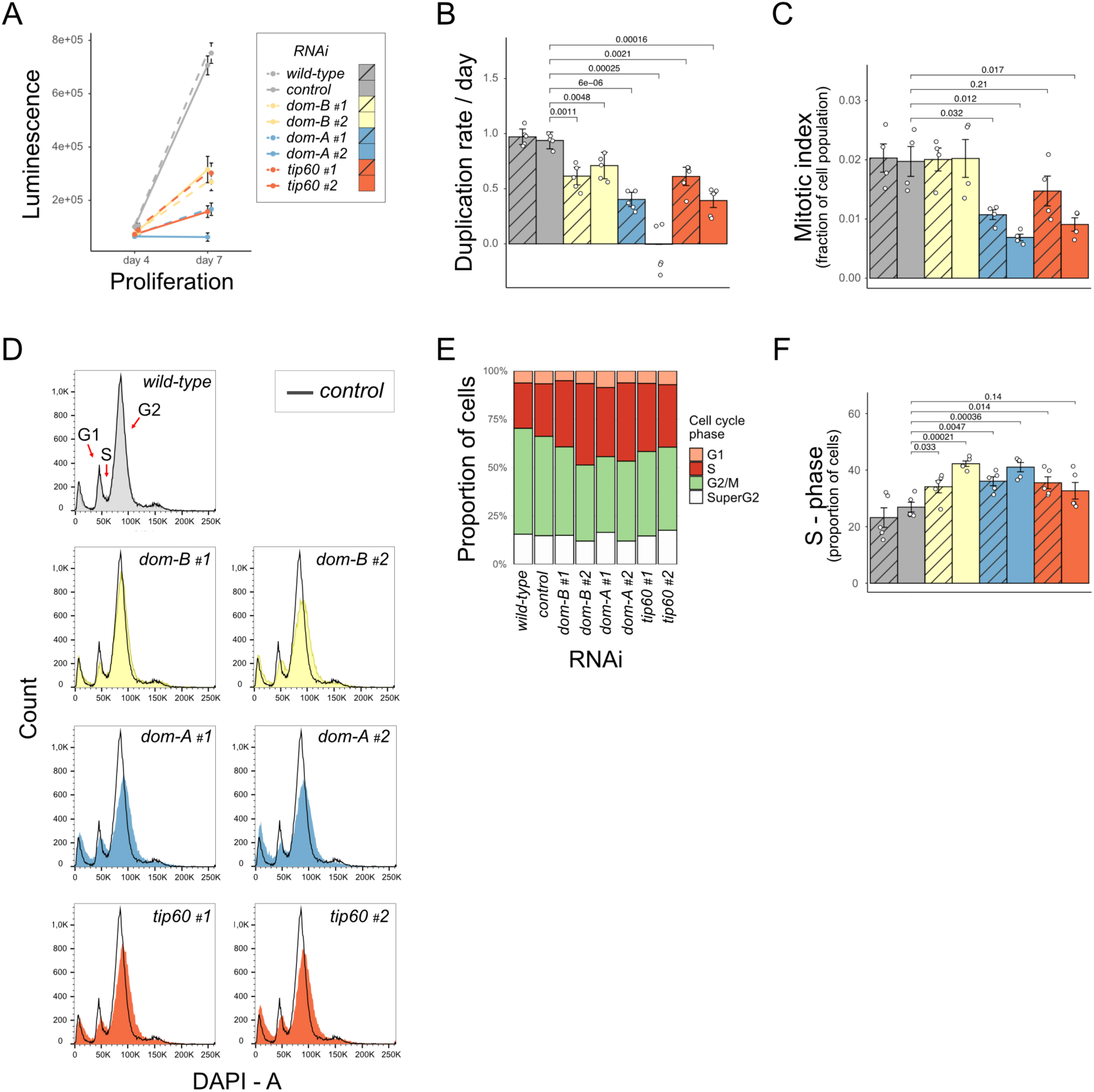
Depletion of Dom-A/Tip60 leads to cell cycle arrest in Kc167 cells. A) Viability of Kc167 cells upon RNA interference (RNAi) with *tip60*, *dom-A*, *dom-B* expression. Control cells were treated with RNAi against *gst*. Viability was measured using the CellTiter-Glo assay on day 4 and day 7 of RNAi treatment. Error bars: standard error of the mean (SEM) of 6 biological replicates for day 4 and 5 biological replicates for day 7. Different dsRNA sequences targeting the same transcript are indicated by #1 and #2. B) The data in (A) were replotted to reflect the duplication rate of the cells, as indicated. C) Mitotic index of cells (measured by H3S10ph staining) depleted of Tip60, Dom-A, Dom-B or control cells. Error bars: SEM of 4 biological replicates. D) The cell cycle profile of wild-type and control cells or cells lacking Tip60, Dom-A, Dom-B was determined by DAPI (4′,6-diamidino-2-phenylindole) staining followed by flow cytometry. E) Cell cycle phase distribution of the cell populations analyzed in (D). F) Quantification of S-phase in (D). Error bars: SEM of 5 biological replicates.

To quantify the impact on cell proliferation, we calculated duplication rates from the CellTiter-Glo measurements (Figure 1B). *Tip60* and *dom-A* knockdown significantly impaired duplication rates relative to controls, documenting the importance of the DOM-A/TIP60 complex in sustaining cell proliferation. Furthermore, RNAi against *tip60* and *dom-A* resulted in a pronounced reduction in the mitotic index, measured by H3S10 phosphorylation (Figure 1C). In contrast, depletion of *dom-B* did not have a detectable effect in the mitotic index, underscoring the distinct functions of the DOM-A and DOM-B complexes in cell cycle control.

To assess the cell cycle distribution, we determined DNA content profiles by flow cytometry of RNAi-treated Kc167 cells. As illustrated in Figure 1D-F and Supplementary Figure S1B, both cells depleted of *tip60* or *dom-A* tended to accumulate in S phase with a concomitant reduction of G2-M phase. We also detected an elevated sub-G1 population, indicative of enhanced cell death (Supplementary Figure S1C). These data demonstrate that the DOM-A/TIP60 complex is essential for maintaining efficient cell cycle progression.

### Cells arrested upon Dom-A/Tip60 depletion are not sensitive to irradiation

Given the established role of mammalian TIP60/P400 in the DNA damage response, we tested whether depletion of Dom-A and Tip60 in *Drosophila* Kc167 cells would sensitize cells to exogenous DNA damage. Kc167 cells treated with RNAi targeting *tip60*, *dom-A* or *dom-B* along with the *gst* control were irradiated with X-rays or UV-C on day 4 of RNAi and cell viability was measured on day 7 using the CellTiter-Glo assay (Figure 2A).

**Figure 2.**
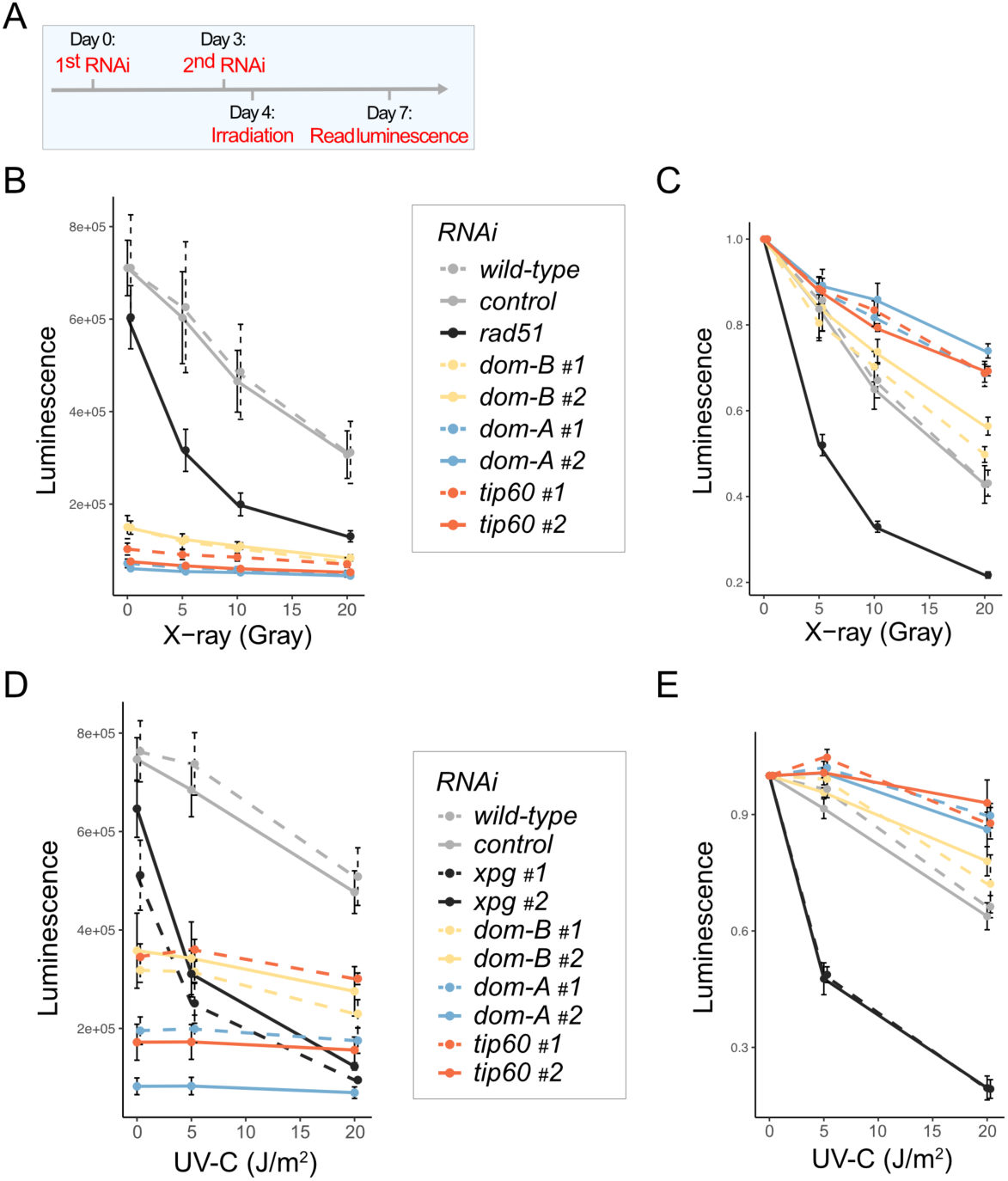
Viable cells upon DOM-A/TIP60 depletion are not sensitive to mutagenic radiation. A) Schematic timeline of the experiment. B) Cells depleted of Tip60, Dom-A, Dom-B or controls were X-irradiated at the indicated dose on day 4 of RNAi treatment. Cell viability relative to non-irradiated conditions was measured on day 7 using the CellTiter-Glo assay. Depletion of dRad51 serves as a control for a factor with known involvement in homologous recombination repair. Error bars: standard error of the mean (SEM) of 3 biological replicates. C) As in (B), but raw intensity values are expressed as a fraction of the non-irradiated condition. D) Cells as in (B) were UV-C-irradiated at the indicated dose on day 4 of RNAi treatment. Cell viability relative to non-irradiated conditions was measured on day 7 using the CellTiter-Glo assay. Depletion of Mus201 serves as a control for a factor with known involvement in UV-C repair. Error bars: SEM of 3 biological replicates. E) As in (D), but raw intensity values are expressed as a fraction of the non-irradiated condition.

Upon X-ray irradiation, we observed a progressive decline in cell viability in control cells. As a reference, depletion of *Drosophila* Rad51 (Spn-A), a key homologous recombination (HR) repair factor, strongly sensitized cells to X-rays without a strong effect on basal proliferation (Figure 2B). At the time of irradiation, the viability of Tip60- and Dom-A-depleted cells was already much compromised, but the remaining cells were remarkably resistant to irradiation. This became apparent when cell viability was displayed relative to the corresponding non-irradiated control (Figure 2C). While *rad51* knockdown significantly sensitized cells to X-rays, depletion of Dom-A or Tip60 did not increase radiation sensitivity. In fact, depletion of Dom-A or Tip60 rendered the remaining cells <u>less</u> sensitive to X-rays compared to control cells. In contrast, *dom-B* depletion affected baseline viability, but did not enhance X-ray sensitivity.

Similar results were obtained following UV-C irradiation. RNAi against *Mus201*, known to be involved in UV repair (*Drosophila xpg*), served as a positive control and showed the expected hypersensitivity (Figure 2D-E). In contrast, neither Tip60-nor Dom-A-depleted cells exhibited increased sensitivity to UV-C when viability was normalized to the unirradiated control (Figure 2E).

These findings indicate that the dominant phenotype following Tip60 or Dom-A depletion is a pronounced cell cycle arrest, which renders the cells relatively resistant to additional cytotoxic effects of DNA damage. These results support the conclusion that the primary function of the DOM-A/TIP60 complex in Kc167 cells is to sustain cell proliferation.

### Deregulation of cell cycle gene expression upon depletion of DOM-A/TIP60

Given the cell cycle arrest phenotype observed upon Tip60 and Dom-A depletion, we next asked whether this effect was reflected at the level of gene expression. To address this, we re-analyzed existing RNA sequencing data from Kc167 cells (15). For comparison, we also included data from Dom-B-depleted cells. Transcriptome profiling revealed that depletion of Tip60 and Dom-A led to highly similar transcriptome signatures, clearly distinct from the effects of either *dom-B* knockdown or control samples (Figure 3A, Supplementary Figure S3A). This illustrates the earlier conclusion (15) that Tip60 and Dom-A function in a complex to regulate gene expression, while Dom-B, the principal H2A.V exchange factor, acts independently.

**Figure 3.**
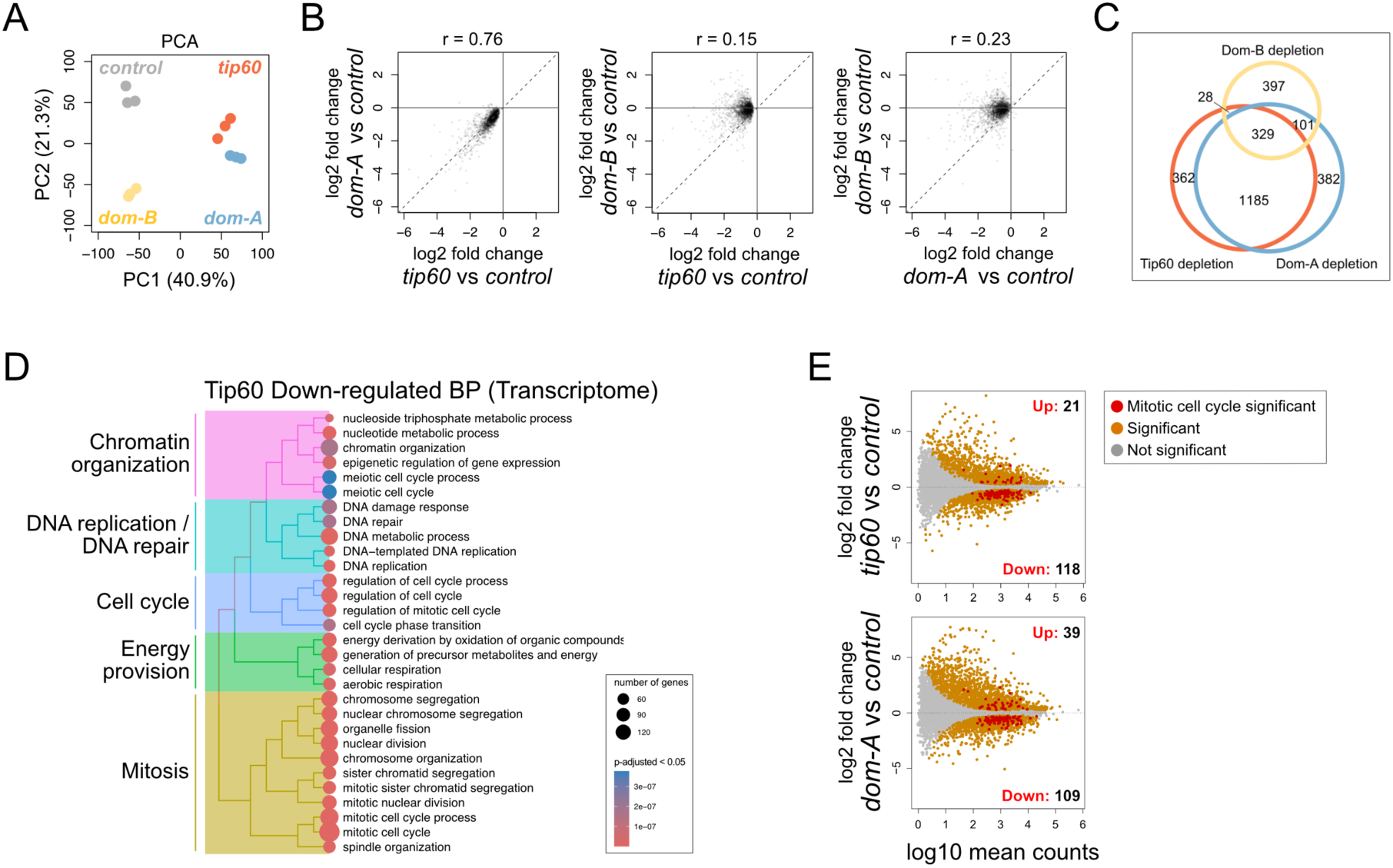
The DOM-A/TIP60 complex promotes transcription of cell cycle genes. A) Principal Component Analysis (PCA) comparing transcriptome profiles derived from Kc167 cells depleted of Tip60, Dom-A or Dom-B. Data from 3 biological replicates were used. The percentage of variance associated with PC1 and PC2 is indicated in parenthesis. B) Scatter plots comparing the log2 fold-changes in expression of *tip60* down-regulated genes (filtered by p-adjusted cut-off < 0.05; N = 1904) across the depleted cell lines shown in (A). The Spearman correlation coefficient (r) is shown at the top of the plot. C) Venn diagram indicating the overlap of down-regulated genes of the data in (B). D) Dendrogram showing the top 30 significantly enriched biological process (BP) GO terms from ORA of genes downregulated upon Tip60 depletion, filtered by p-adjusted cut-off < 0.05. The color of the circles indicates enrichment and their size the number of genes annotated with the corresponding pathway. GO terms were clustered based on semantic similarity and interpretation summaries are given to the left. E) MA plot comparing log2 fold-change to mean RNA-seq counts, derived from three independent experiments between control Kc167 cells and cells depleted of either Tip60 or Dom-A. Differentially expressed genes within the indicated gene group are shown in red (p-adjusted cut-off < 0.05), differentially expressed genes but not within the corresponding gene group are shown in orange, and non-differentially expressed genes and not within the gene group are shown in grey. Numbers indicate up-or down-regulated mitotic cell cycle significant genes (red group).

Indeed, the significantly downregulated genes upon Tip60 depletion (N = 1904, p-adjusted < 0.05) are similarly affected by Dom-A depletion (Figure 3B). *Tip60* and *dom-A* knockdowns showed a strong positive correlation, which was not the case for a *dom-B* knockdown. There was a substantial overlap in downregulated genes of Tip60 and Dom-A depletion, with a more limited intersection with *dom-B* (Figure 3C).

Gene Ontology (GO) analysis of the genes with lowered expression in the absence of Tip60 reveals a significant enrichment for categories related to proliferation, cell cycle progression and energy metabolism (Figure 3D; for GO with higher expression, see Supplementary Figure S3B). Notably, GO categories associated with mitotic processes were predominantly downregulated, suggesting a broad role for Tip60 in sustaining proliferative transcription programs. Complementary, GO analysis of Dom-A-regulated genes produced similar results, with enrichment for mitosis and nuclear division pathways (Supplementary Figure S3C). Comparison of RNA-seq log2 fold-changes between control and Tip60-or Dom-A-depleted cells confirms the reduction of proliferation-associated transcripts, including many with roles in the mitotic cell cycle (Figure 3E). In contrast, depletion of Dom-B only modestly impacted mitotic cell cycle genes (Supplementary Figure S3D).

### Proteome-wide alteration in cell cycle regulators following TIP60 depletion

To complement the transcriptomic analysis, we determined the changes in nuclear proteome upon Tip60 depletion. For comparison, we carried out a parallel analysis with cells depleted of the MYST-family acetyltransferase chameau (Chm, KAT7). We processed four biological replicates for each condition and identified more than 3000 proteins (Supplementary Tables 2 and 3 represent Class1 and 2, respectively). Proteomic profiles showed distinct clustering of Tip60-depleted replicates, while cells with reduced Chm levels were more similar to the control (Figure 4A). Analysis of proteome profiles revealed widespread differential expression of nuclear proteins in Tip60-depleted cells, whereas changes following Chm depletion were modest by comparison. (Figure 4B). Global analysis of significantly altered nuclear proteins further emphasized the widespread impact of Tip60 depletion (Supplementary Figure S4A).

**Figure 4.**
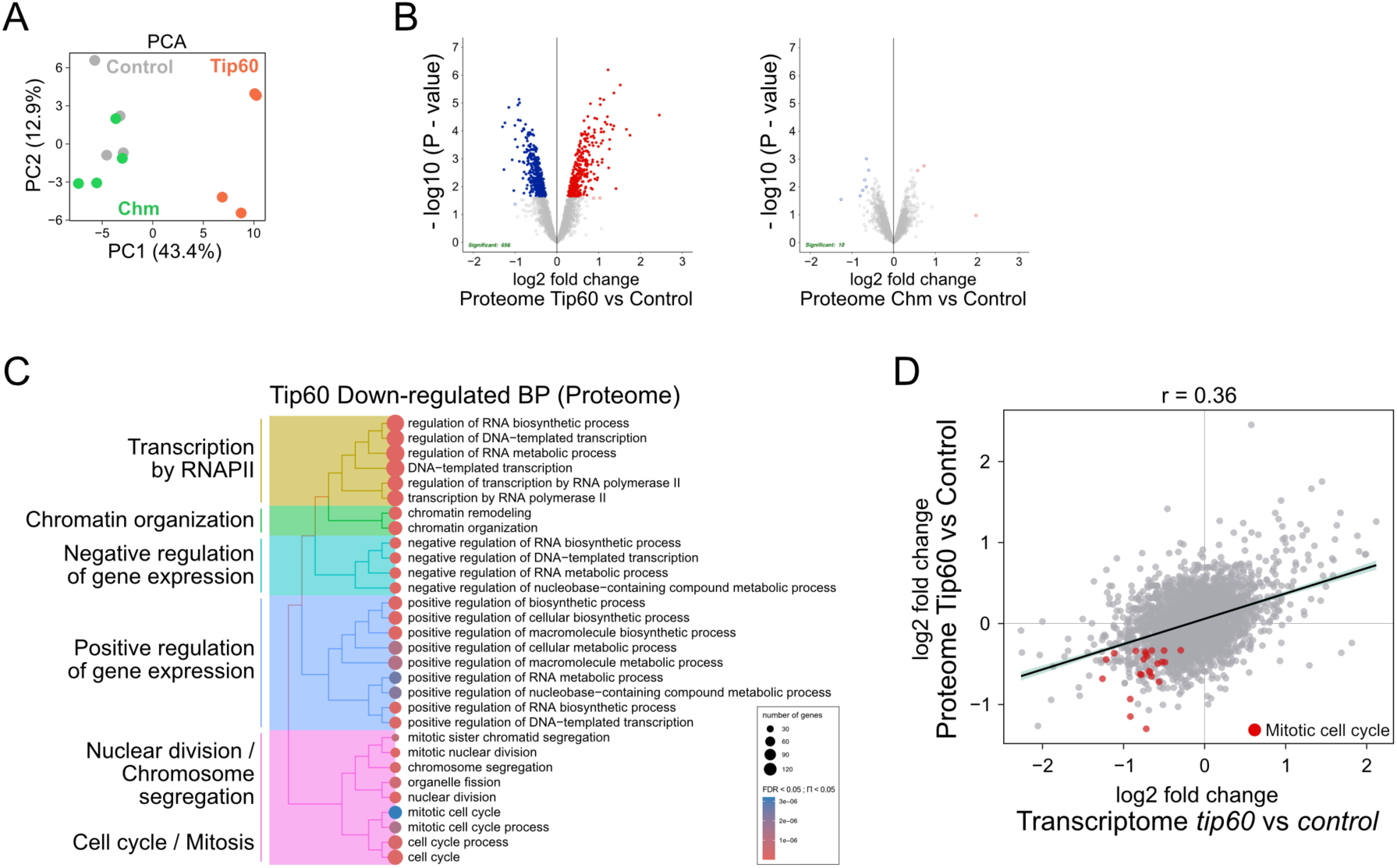
Deregulation of cell cycle proteins upon depletion of Tip60. A) PCA comparing nuclear proteome profiles derived from Kc167 cells depleted of Tip60, Chm or Control by RNAi. Data from 4 biological replicates were merged. PC1 and PC2 are shown. Percentage of variance is indicated in parenthesis. B) Volcano plot showing log2 fold-change in the x-axis and log10 P-value in the y-axis for ‘Class 2’ proteins for Tip60-depleted cells vs control (Left), and Chm-depleted cells vs control (Right). Filled circles indicate FDR < 0.05 and unfilled circles indicate Π < 0.05. Color of the circles indicates the nature of differential regulation (red: up; blue: down). C) Dendrogram showing the top 30 significantly downregulated Biological Process (BP) GO terms for combined Class 1+2 proteins from ORA of the Tip60-depleted proteome, filtered by FDR < 0.05 and Π < 0.05. Color of the circles indicates enrichment and their size indicates number of genes annotated with the corresponding pathway. GO terms were clustered based on semantic similarity and the terms that were represented the most within a cluster were mentioned. D) Correlation comparing the log2 fold-change in expression between Tip60-depleted proteome and transcriptome (N = 2881). Common genes between proteome and transcriptome are shown in grey, and genes within the gene group ‘mitotic cell cycle’ filtered for p-adjusted < 0.01 (transcriptome) and FDR < 0.05 and Π < 0.05 (proteome) are shown in red. The Spearman correlation coefficient (r) is shown at the top of the plot.

GO analysis of the proteins that were reduced upon Tip60 depletion mirrored the transcriptomic findings, with top enriched categories relating to cell cycle, nuclear division, and chromatin regulation (Figure 4C). The log2 fold-changes in RNA and protein expression for genes present in both datasets were correlated (Figure 4D), and although the correlation was moderate (r = 0.36), it suggested that protein levels were regulated by both, transcription and protein stability.

We also examined whether depletion of Tip60 altered the abundance of its own complex components. Label-free quantification (LFQ) revealed that loss of Tip60 led to a reduction in Ing3 and Eaf6, two subunits of the TIP60 core module, while Chm depletion did not (Supplementary Figure S4B).

Collectively, these data reinforce the view that the DOM-A/TIP60 complex acts as a critical regulator of essential proliferation genes in Kc167 cells.

### Tip60 preferentially binds a subset of active promoters and correlates with H2A.V acetylation

The acetyltransferase Tip60 has prominent substrates in chromatin, notably the N-termini of histones H4 and H2A.V. In particular, H2A.VK4/7 at the first nucleosome downstream of the transcription start site (TSS), the ‘+1’ nucleosome, is a hallmark of active promoters.

To investigate the genomic localization of Tip60 along with its substrate H2A.V and acetylated H2A.V (H2A.Vac), we performed CUT&RUN profiling in Kc167 cells. Hierarchical clustering of CUT&RUN data of biological replicates documents the reproducibility of the profiles (Supplementary Figure S5A). Visualization of the Tip60 peaks in the heatmaps reveals two clusters of Tip60 binding sites in relation to the H2A.V localization (Figure 5A). In cluster 1, Tip60 binding colocalizes with (acetylated) H2A.V, while cluster 2 shows Tip60 at sites that lack H2A.V. The annotation of genomic features reveals that cluster 1 binding sites are predominantly located at promoters, whereas cluster 2 sites were more often found within introns and intergenic regions (Figure 5B).

**Figure 5.**
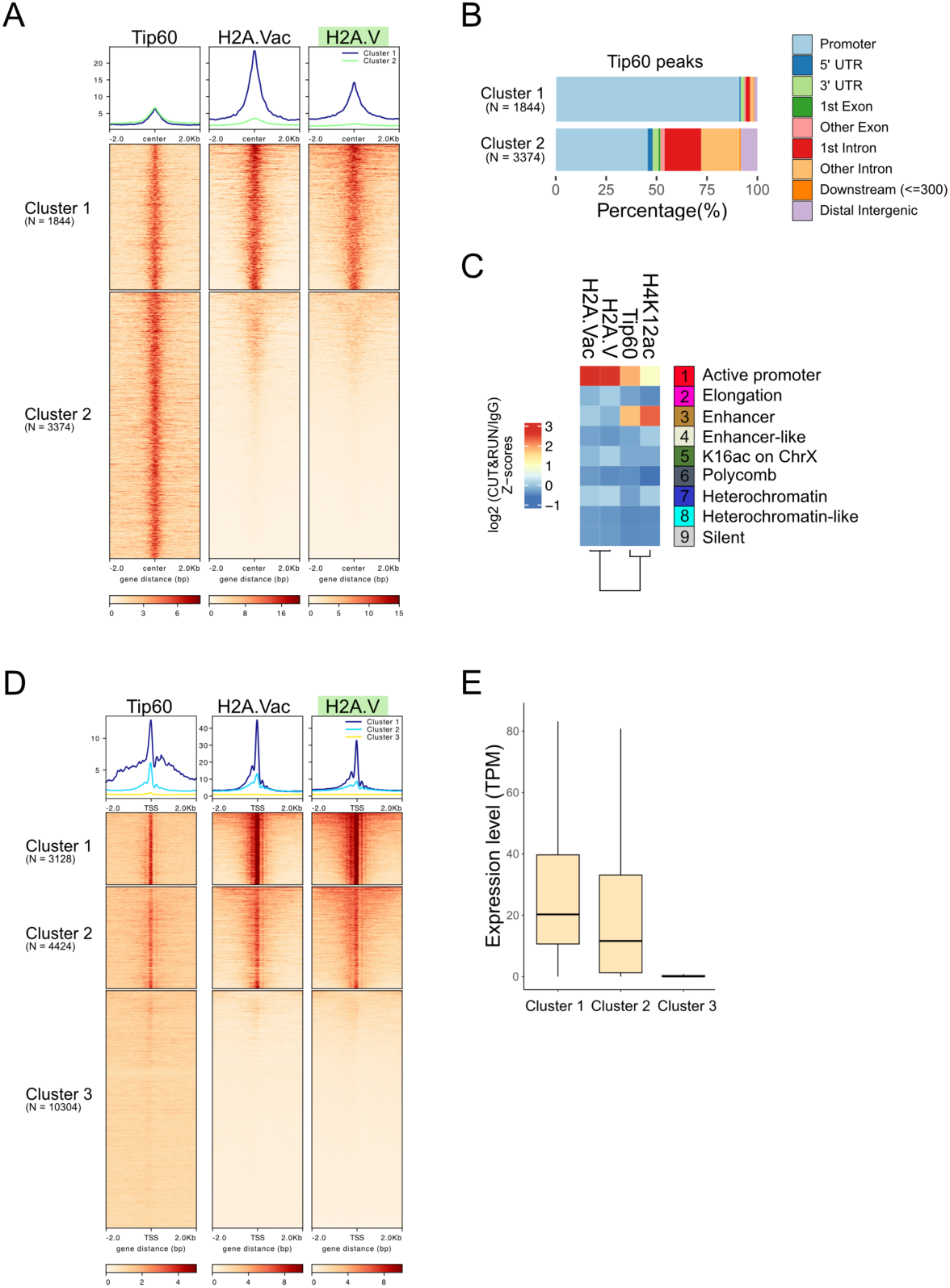
TIP60 binding at active promoters correlates with acetylation of H2A.V. A) The genomic binding sites of Tip60 (peaks from CUT&RUN analysis) were clustered as a function of H2A.V localization. Protein enrichment is normalized to the corresponding control and presented as cumulative profiles (top) and heat maps (bottom). Coverage windows of 4 kb around the peaks are shown and values correspond to the average of 3 biological replicates. Loci are sorted into 2 clusters (k-means) based on H2A.V (highlighted green). H2A.V localization coincides with the acetylated form (H2A.Vac). B) Genomic annotation of Tip60 peaks based on the two clusters in (A). C) Chromatin state enrichment analysis using a nine-state ChromHMM (modENCODE) (42) of 3 biological replicates for Tip60, H2A.Vac, H2A.V and one replicate for H4K12ac CUT&RUN profiles. Hierarchical clustering of profiles reveals distinct patterns of enrichment across chromatin states. D) Enrichment of Tip60, H2A.Vac and H2A.V by CUT&RUN on transcription start sites (TSS) of 17856 *Drosophila* genes, by cumulative profiles (top) and heatmaps (bottom). Coverage windows of 4 kb around the TSS are shown and the mean is calculated for each column. Loci are sorted into three clusters (k-means) based on H2A.V (highlighted green). The coverage of each replicate was normalized to its corresponding IgG control (for H2A.V and H2A.Vac) or pre-immune serum (for Tip60), and the mean signal of 3 biological replicates was calculated. E) Comparison of the loci grouped in the three clusters in (D) with the RNA-seq expression level (TPM) of control cells.

Chromatin state enrichment analysis using a nine-state ChromHMM model (42) further refined these observations (Figure 5C). Hierarchical clustering revealed that Tip60 predominantly occupies active promoter and enhancer regions, while H2A.Vac was restricted to promoters. Notably, consistent with previous findings (43), H2A.V was largely absent from enhancer regions, supporting the view that TIP60-mediated H2A.V acetylation is largely confined to promoters. Interestingly, H4K12 acetylation was found to be predominantly enriched at enhancer regions, with comparatively lower levels at promoters, suggesting that TIP60-mediated H4K12 acetylation is largely enhancer-specific. This pattern of H4K12 acetylation was further validated by ChIP-seq assay (Supplementary Figure S5B). These findings point to an interesting locus-specific functional diversification according to which at H2A.V acetylation is the *modus operandi* at promoters, whereas H4K12 acetylation is favored at enhancers.

Changing the perspective, we monitored the association of Tip60 at the transcription start sites (TSS) of 17,856 annotated *Drosophila* genes (Figure 5D). CUT&RUN profiles show that Tip60 localizes specifically to the subset of promoters (clusters 1 and 2) characterized by H2A.V enrichment. The patterns resolve the nucleosomes around the TSS, with highest H2A.V enrichment at the ‘+1’-nucleosome. The H2A.V acetylation pattern follows that of H2A.V. This feature appears to have functional significance, since Tip60 and H2A.Vac occupancy was positively correlated with steady-state gene expression (TPM) in control Kc167 cells (Figure 5E).

Together, these findings suggest that the DOM-A/TIP60 complex targets a distinct cohort of active promoters where H2A.V acetylation correlates with transcriptional activation, while Tip60 binding at enhancers may reflect functions independent of H2A.Vac.

### Depletion of Tip60 leads to a global reduction of H2A.V acetylation at promoters

The localization of Tip60 correlated well with H2A.V acetylation at promoters. To explore causality, we depleted Tip60 and analyzed the levels of residual Tip60 and H2A.Vac by immunoblotting of chromatin-enriched fractions. This revealed a pronounced reduction in acetylated H2A.V levels (Supplementary Figure S6A). CUT&RUN profiling showed that Tip60 signals were effectively lost at TSS genome-wide, as expected (Figure 6A). This correlated with a substantial reduction in H2A.Vac at promoters, indicating that Tip60 is indeed the primary enzyme responsible for acetylating H2A.V at these sites. A genome-wide browser view further illustrated the widespread reduction of both Tip60 binding and H2A.Vac upon Tip60 depletion across *Drosophila* chromosomes (Figure 6B).

**Figure 6.**
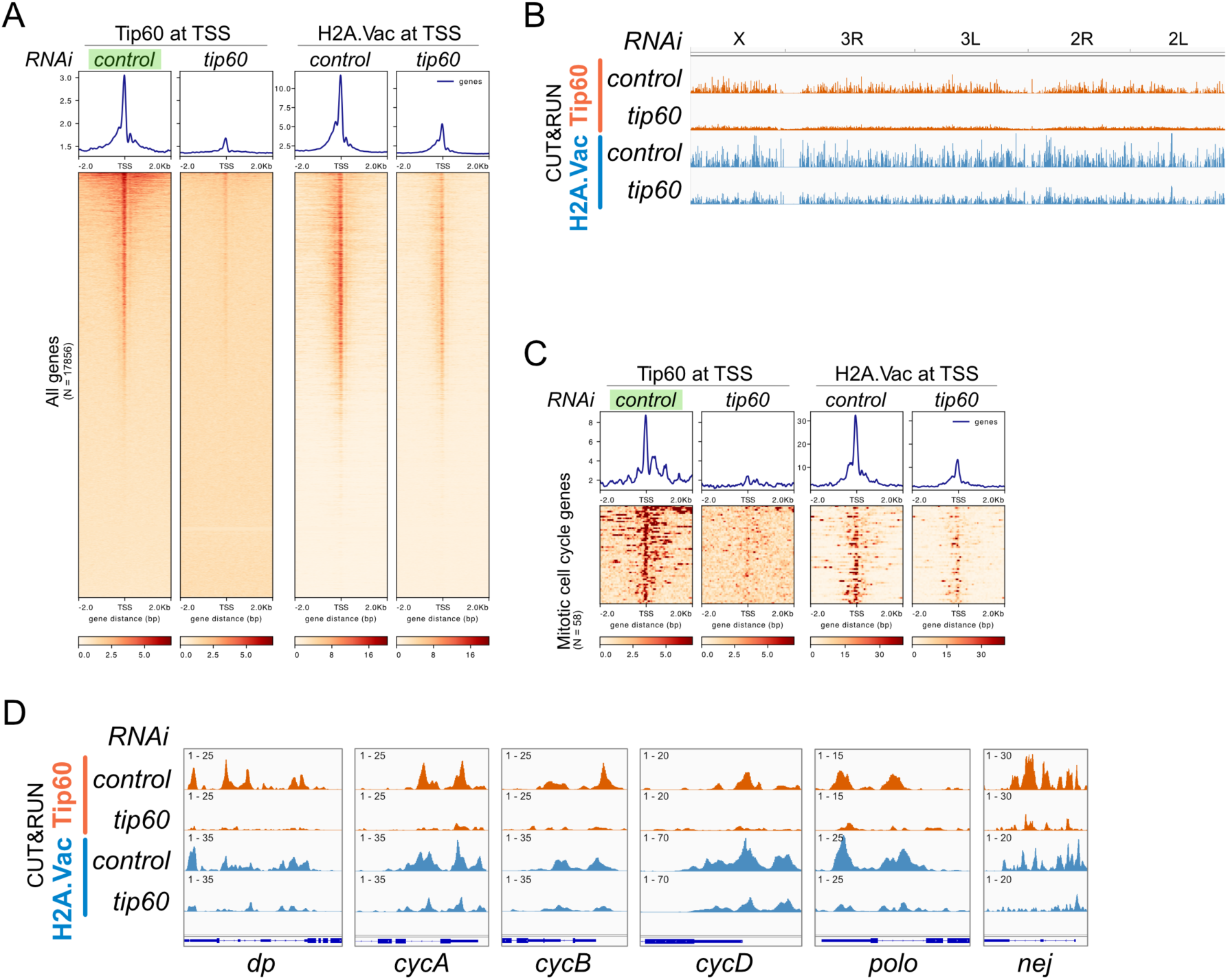
Depletion of TIP60 leads to reduced H2A.Vac. A) Enrichment of Tip60 and H2A.Vac by CUT&RUN on TSSs (N = 17856) in Kc167 cells depleted of Tip60 (*tip60*) relative to control, illustrated by cumulative profiles (top) and heatmaps (bottom). Coverage windows of 4 kb around the TSS are selected and the mean is calculated for each column. Sorting is according to the Tip60 signal in control cells (highlighted green). The coverage of each replicate was normalized to its corresponding IgG control (for H2A.V and H2A.Vac) or pre-immune serum (for Tip60), and the mean signal of 3 biological replicates was calculated. B) Genome browser tracks of Tip60 and H2A.Vac CUT&RUN in control cells or cell RNAi-depleted of Tip60 (*tip60*) across the *Drosophila melanogaster* chromosomes. C) Enrichment of Tip60 and H2A.Vac around the TSS of genes with GO annotation ‘mitotic cell cycle’ selected for binding of Tip60 and reduced expression upon Tip60-depletion (q-adj < 0.05). The display is as in (A). D) Genome browser tracks of Tip60 and H2A.Vac CUT&RUN in control cells or upon RNAi depletion of Tip60 (*tip60*) at promoters of the genes *Dp*, *CycA*, *CycB*, *CycD*, *polo* and *nej*.

Focusing on a subset of genes associated with the GO term ‘mitotic cell cycle’ (Supplementary Figure S6B), of which many showed reduced expression in Tip60-depleted cells, we observed marked losses of Tip60 and H2A.Vac at corresponding TSS (Figure 6C). Among them are 6 subunits of the replicative helicase complex (Supplementary (Figure S6B). Figure 6D exemplifies these general statements with genome browser tracks for relevant genes, such as genes encoding a transcription factor E2F subunit (*Dp)*, various cyclins (*CycA*, *CycB*, *CycD)*, polo kinase (*Polo*), and *Drosophila* CBP (*nej*).

We also found that Tip60 and H2A.Vac are prominently enriched at many promoters of genes with GO term ‘DNA repair’. Here the H2A.Vac signals were also substantially diminished following Tip60 depletion (Supplementary Figure S6C, D).

Together, these findings demonstrate that Tip60 is required for H2A.V acetylation at promoters on a genome-wide scale, and that this modification is particularly important for the expression of genes driving both cell cycle progression and genome integrity maintenance in *Drosophila* cells.

### Depletion of Tip60 leads to widespread alterations in the nuclear acetylome

The correlation between the transcriptome and proteome changes upon Tip60 depletion were only modest (Figure 4D), suggesting an additional layer of regulation. Because protein stability can be affected by acetylation through competition with ubiquitylation at relevant lysines (44), we considered contributions of Tip60-dependent acetylation. Yeast and mammalian TIP60 are known to acetylate proteins in addition to histones (45–48). To explore the broader acetylation landscape governed by Tip60 in *Drosophila*, we profiled the nuclear acetylome of Kc167 cells by mass spectrometry following RNAi-mediated depletion of Tip60 relative to control. As a further reference, we determined the acetylome changes upon depletion of Chm.

We processed four biological replicates for each condition and identified more than 430 acetylated peptides, which correspond to more than 330 acetylated proteins, some of which were acetylated at multiple sites. One class of peptides ‘Class 1’ was identified robustly in most or all control conditions, but not at all or only in one or two Tip60 depletion replicates. These are the strongest candidates for Tip60-dependent acetylation sites. Because multiple values were missing, imputation of missing values was not warranted. As a result, we do not have statistical information about these ‘Class 1’ peptides and they have also not been normalized to the corresponding proteome values (Supplementary Table 4). The absence of a peptide may to some extent be explained by the absence of the protein, for example due to the downregulation of gene expression discussed earlier. ‘Class 2’ peptides were identified and quantified in most of the samples and missing values were imputed to obtain quantitative data, which were normalized to the ‘input’ proteome (Supplementary Table 5).

The acetylome profiles (all peptides) separated the Tip60-depleted samples clearly from control cells, accounting for the largest proportion of variance (Figure 7A). This indicates that loss of Tip60 induces specific changes in the nuclear acetylome. Tip60 and Chm share some substrates (Figure 7B), which likely accounts for the partial overlap observed between the sample groups. Approximately 65% of the Tip60-dependent acetylation events were specific to Tip60 depletion. Likewise, approximately half of the acetylation changes upon Chm depletion appeared specific to this KAT, demonstrating the distinct substrate specificities of these two MYST family acetyltransferases.

**Figure 7.**
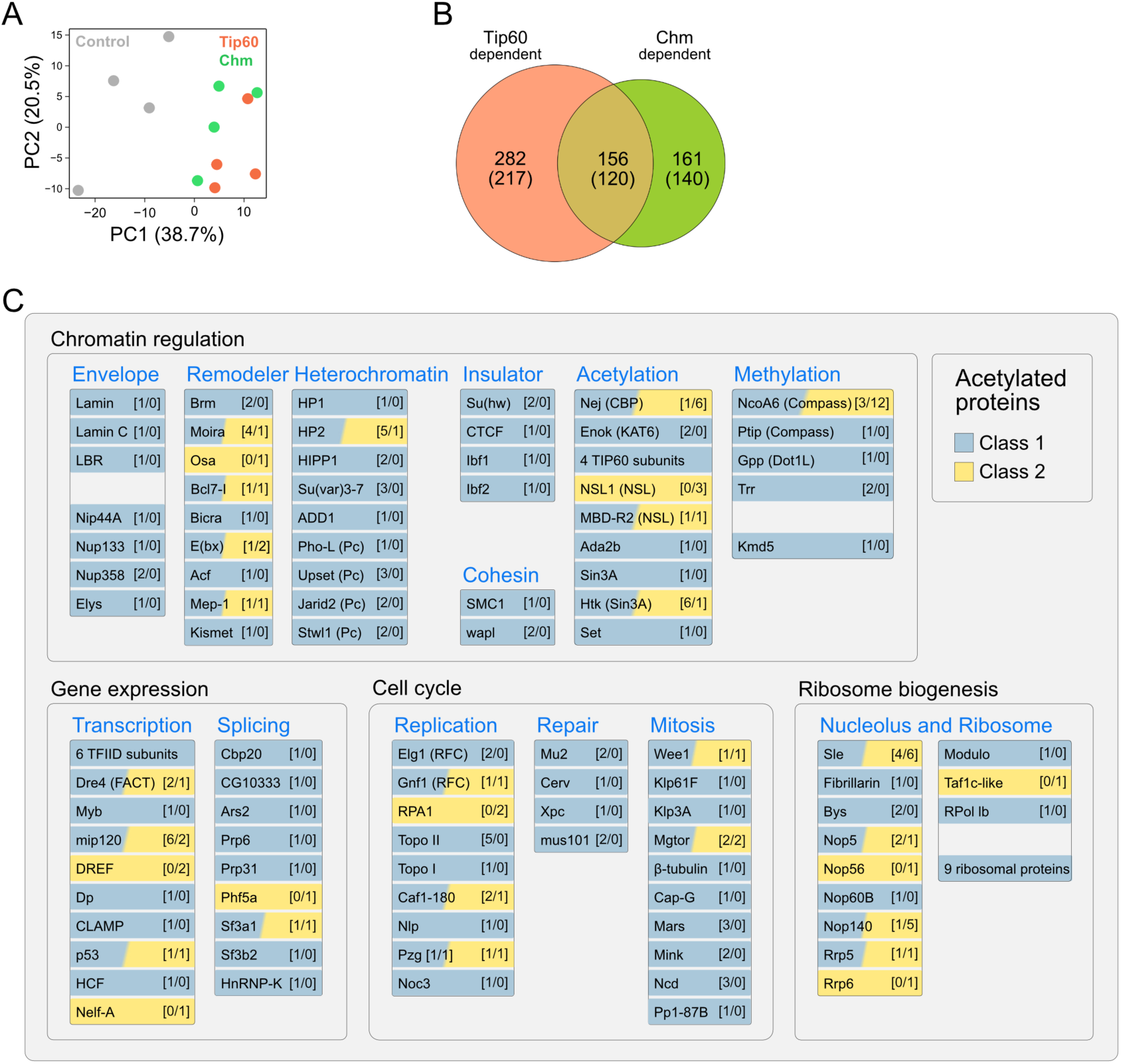
Depletion of Tip60 leads to numerous acetylome changes. A) Principal Component Analysis (PCA) of nuclear acetylome profiles from Kc167 cells following RNAi-depletion of Tip60, Chm, or Control. Data from 4 biological replicates were combined. The percentage variance associated with PC1 and PC2 is indicated in parentheses. B) Venn diagram showing the overlap between the acetylated peptide targets (‘Class 1’ and ‘Class 2’) that are regulated by Tip60 and Chm. The number in parenthesis indicate the number of proteins. The ‘Class 2’ peptides were filtered using thresholds of FDR < 0.05 and Π < 0.05. C) Visualization of selected acetylated protein targets regulated by Tip60 (for full list see Supplementary Tables 4 and 5). Proteins categorized from ‘Class 1’ peptides are shown in blue, while ‘Class 2’ proteins are displayed in yellow. The number in brackets indicate the number of different acetylated peptides identified in ‘Class 1’ (on the left) or ‘Class 2’ (on the right).

Focusing on Tip60, widespread changes in differentially acetylated ‘Class 2’ peptides were observed following Tip60 loss (Supplementary Figure S7A, B). While acetylation gains are unquestionable due to indirect effects (see below), peptides that lose acetylation upon Tip60 depletion qualify for direct targets. GO analysis of those proteins reveals a pronounced enrichment for categories related to transcription, chromatin organization and RNA/DNA metabolism (Supplementary Figure S7C). Related GO categories were enriched following Chm depletion (Supplementary Figure S7A, B, D).

Inspection of the lists of Tip60-dependent ‘Class 1’ peptides or significantly changed ‘Class 2’ peptides yields a remarkable set of proteins involved in replication and mitosis, as well as gene expression and chromatin regulation with relevance to cell growth and proliferation. Figure 7C lists selected to illustrate the potential of acetylation-based regulation. We find four different TFIID subunits, a FACT subunit (Dre4), several sequence-specific activating transcription factors (TFs), among them p53, myb, DREF, the pioneer TF CLAMP and HCF, as well as repressors of transcription initiation (tramtrack, ttk) and elongation (Nelf-A). Furthermore, a substantial number of factors involved in transcript splicing and processing are found. The functional term ‘cell cycle’ lists several proteins involved in replication, among them both topoisomerases, the histone chaperone subunit Caf1-180, two RFC subunits and RPA, as well as several proteins are involved in mitosis. Surprisingly, a large number of nucleolar proteins and ribosome subunits are found acetylated.

Among the epigenetic regulators that are candidates for Tip60-dependent acetylation are proteins of nucleosome remodeling complexes, chromosomal insulators, components of the nuclear lamina and nuclear pores, proteins involved in forming constitutive and facultative (polycomb-dependent) heterochromatin, protein methyltransferases and demethylases, as well as proteins involved in acetylation/ deacetylation reactions.

In summary, these results demonstrate that Tip60 brings about a unique nuclear acetylome signature in *Drosophila* proliferating cells, involving numerous proteins involved in gene expression, chromatin regulation and cell cycle control.

## Discussion

Our comprehensive analysis of the consequences of Tip60 depletion in a *Drosophila* cell model reveals the DOM-A/TIP60 complex as an essential regulator of proliferation and cell cycle progression. Depletion of the effector acetyltransferase Tip60 or its partner Dom-A causes marked proliferation defects, a pronounced S-G2/M cell cycle arrest, and broad transcriptional and proteomic changes with relevance for cell growth and proliferation. We hypothesize that Tip60, through the compendium of its prominent substrates, plays a central role in coordinating cellular functions to assure a smooth progression through the cell cycle. Our findings are in line with recent genetic and molecular studies in mammals, where TIP60 deletion or depletion caused cell cycle arrest with failure to proceed to metaphase (17,49), suggesting they may have more general relevance.

Histones are prominent substrates of Tip60 and their acetylation at promoters immediately suggests a mechanism for transcriptional regulation. In addition, our description of the Tip60-dependent acetylome in *Drosophila* widens the spectrum of potential, non-exclusive mechanisms of proliferation control.

### Tip60 acetylates H2A.V at the promoter of cell cycle genes

One evolutionary conserved mechanism through which Tip60 regulates transcription is the acetylation of histones H4 at lysine 12 and of the H2A variant H2A.Z (H2A.V in *Drosophila*). The acetylation of H2A.Z at the first nucleosome downstream of the transcription start site characterizes active promoters (10). We found that Tip60 binds to active promoters and promotes acetylation of H2A.V (H2A.Vac). Depletion of the KAT leads to reduced transcription of critical genes involved in cell cycle regulation, such as cyclins A, B and D, polo kinase and the E2F subunit dp, which explains the proliferation arrest. Interestingly, our data suggest that Tip60-dependent H4K12 acetylation (15,50) is prominent at enhancers, where H2A.Vac is not enriched.

### Beyond gene regulation: implications of the Tip60-dependent acetylome

A recent cell cycle-resolved acetylome study in HeLa cells revealed the complexity of acetylation in proteins other than histones and their changes during the cell cycle (47). For *Drosophila*, such a comprehensive analysis had been lacking. We focused on the nuclear acetylome of asynchronous cells, and monitored changes upon Tip60 depletion. As a reference for selectivity, we also determined the acetylome catalyzed by the MYST family acetyltransferase Chameau (KAT7) (51), which, like Tip60, is known to acetylate H4K12. While Chm affects proteins involved in intermediary metabolism (52,53), Tip60 substrates relate to gene expression, chromatin biology and the cell cycle. At this point, we have no evidence that any of these acetylations is functional, however, the nature of the acetylated proteins, including powerful regulators of transcription and chromatin structure, replication and mitosis, nucleolus organization and ribosome assembly, is striking. In the following, we will speculate about the regulatory potential of the Tip60 acetylome.

### Acetylation of transcription factors

The TIP60-dependent acetylome includes factors generally involved in transcription initiation (6 subunits of the general transcription factor TFIID), elongation (the FACT subunit Dre4, Nelf-A, topoisomerase I), as well as in co-transcriptional splicing/ RNA processing. The latter observation resonates with recent observations of functional interactions of Tip60 with the splicing machinery (54). It has been reported that Tip60 binds directly to nascent RNA (55) and this may well go along with acetylation of splicing factors.

TIP60 has been described as a transcriptional co-activator complex that mediates the action of specific transcription factors (TFs) involved in cell cycle progression, such as c-Myc, E2F1and p53 (6,17). In flies, the DOM-A/TIP60 complex directly interacts and functionally cooperates with the transcription factor Myc in neuroblast maintenance (56). We find several specific TFs acetylated that are known to regulate S-phase genes or to mediate signals for cell cycle progression. For example, DREF is involved in mediating signals from TOR, JUNK and EGFR pathways (57). Among the high-confidence TIP60-dependent acetylation targets are several subunits of DREAM, a complex that regulates hundreds of genes involved in cell cycle decisions (58): the TF Myb and its interaction partner Mip120, as well as the E2F-subunit Dp. Others have suggested roles for Tip60 in regulation of E2F target genes and described genetic interactions with DREAM/MMB components.

### Acetylation of epigenetic regulators

Tip60-dependent acetylation of constituents of the nuclear lamina, nuclear pores, constitutive as well as facultative heterochromatin, chromosomal insulators and cohesin may impact nuclear architecture. With four subunits of the developmental transcription regulator polycomb and three subunits of the COMPASS methyltransferase, two epigenetic antagonists of cell fates and differentiation are found multiply acetylated (59). Remarkably, we found the COMPASS subunit NcoA6, which mediates the effect of the hippo pathway, acetylated at 15 different sites. Acetylations are further found in five distinct nucleosome remodeling complexes, including 5 BRM (brahma) subunits.

Several of the proteins acetylated in a Tip60-dependent manner are involved in acetylation reactions themselves. The most remarkable finding is that *Drosophila* KAT3 (dCBP, encoded by the gene nejire), a central regulator of cell proliferation (60), is acetylated at 7 different sites. Further Tip60 substrates in this category are the acetyltransferase KAT6 (enoki) (61), the KAT8-containing NSL complex (62), as well as components of the Sin3A and INHAT histone deacetylase complexes. If the activity of these enzymes was regulated by acetylation, some of the Tip60-dependent effects may be indirect.

These observations lead us to speculate about the existence of a highly connected regulatory network involving protein acetylation, of which Tip60 is a central player. Acetylation appears to be particularly suited to coordinate critical decision of cell growth and division, because it is intimately connected to the nutritional and energetic state of a cell through the central metabolite acetyl-CoA.

### Nucleolus and ribosomes

Cell proliferation is tightly connected to cell growth, which in turn requires scaling of protein synthesis capacity. Cellular stress is signaled to the nucleolus to dampen the energy-consuming biogenesis of ribosomes (and hence translation), a process that is closely associated with p53-dependent cell cycle arrest (63). Remarkably, we do not only find p53 among the Tip60-dependent acetylome, but 12 proteins involved in nucleolus organization and ribosome biogenesis, including a subunit of RNA polymerase I. Furthermore, 9 ribosomal proteins are acetylated in a Tip60-dependent manner. It is tempting to speculate that acetylation could be used to signal forthcoming conditions for protein synthesis, such as absence of stress and energy availability.

### Progression through S-phase, replication stress and mitosis

In line with the hypothesis that widespread, Tip60-dependent acetylation could promote cell cycle progression, we observed numerous factors involved in replication among the acetylome, including both topoisomerases, two histone chaperones, RPA1, as well as two subunits of the evolutionary conserved replication factor C, Elg1 and Gnf1. Elg1 is a component of the RFC-like complex, which plays a key role in unloading PCNA from chromatin after DNA replication or during replication stress (64), an essential step for maintaining genome stability and enabling cell cycle progression. Tip60 may indeed be involved in modulating replication fork stability. Buttitta and colleagues concluded from their work on mutant imaginal disks that one function of Tip60 *in vivo* is to suppress an endogenous damage response to allow progression through S and G2 phases (65). Notably, we identified SHPRH as a DOM-A interactor, the fly ortholog of yeast Rad5 (unpublished), which reinforces the idea that DOM-A/TIP60 may play a role in post-replicative repair or in resolving replication stress.

Tip60 depletion led to cell cycle arrest, but the remaining viable cells were not particularly sensitive to either X-or UV-irradiation and even seemed to be less sensitive than control cells. This finding is reminiscent of the situation in fission yeast, where the Esa1-Vid21 complex, a functional ortholog of NuA4/TIP60, primarily supports repair of collapsed replication forks, rather than chromosomal breaks (66). This complements emerging models proposing that histone acetylation by TIP60 may modulate replication fork stability and chromatin accessibility during S-phase, rather than operating as a primary responder to acute genotoxic stress (67).

Tip60 also acetylates Mxc, a known substrate of the Cyclin E/Cdk2 kinase complex and a scaffold protein necessary for assembling the histone locus body. This nuclear compartment coordinates transcription and processing of replication-dependent histone mRNAs during S-phase (68). Stalled replication forks could be signaled via Tip60 to repress histone gene expression.

Finally, we find various components involved in organizing the mitotic spindle and cytokinesis enriched among the Tip60-dependent acetylome. It is tempting to speculate that these acetylations reflect the evolutionary conserved interactions of Tip60 with the mitotic apparatus, which are required for proper mitosis and cytokinesis (69).

### Tip60 – a central promoter of cell proliferation?

We highlighted the above-mentioned factors from the rich acetylome comprising some 330 proteins to illustrate the regulatory potential of the Tip60-dependent acetylome. The acetylome contains numerous unstudied proteins (represented by ‘CG’ gene numbers), whose function needs to be explored. Together, our study provides a rich resource to feed functional studies.

At this point we do not have evidence that the acetylations we describe have functional relevance, but given that lysine acetyltransferases are global regulators of various cellular processes, it seems reasonable to assume their regulatory potential. Many of the proteins acetylated by Tip60 are themselves regulators of gene expression, cell cycle signaling and chromatin structure. We consider that information about the cell’s state that are relevant to proliferation decisions are somehow conveyed to the Dom-A complex and Tip60 then coordinates all relevant processes through two complementary mechanisms involving acetylation. The slower regulation of gene expression through histone acetylation sets the general course of the cell cycle, whereas the fast and reversible acetylation of diverse effector proteins may enable fine-tuning of energy-intensive processes in response to stresses or nutritional shortcomings. Poor growth conditions may be signaled broadly by ubiquitous and abundant deacetylases to reinforce cell cycle checkpoints.

In summary, we propose the DOM-A/TIP60 complex as a master regulator of cell proliferation in *Drosophila*, with functional conservation likely extending to higher eukaryotes.

## Acknowledgements

We would like to thank S. Krause for expressing Tip60 in order to generate polyclonal antibodies. We would also like to thank T. Straub from the BMC Bioinformatics Unit for granting us access to the computational cluster, providing us with advice on bioinformatics and helpful scripts. We thank S. Krebs of LMU LAFUGA for Illumina sequencing. We would also like to thank B. Tast and P. Khosravani from the BMC Flow Cytometry Unit for granting us access to the instruments, training and providing us with advice. We are also grateful to the members of the Becker lab for sharing reagents and providing comments on the manuscript.

## Author contributions

Z.A. (Conceptualization, Data interpretation, Data curation, Formal analysis, Investigation, Validation, Writing—original draft, Writing—review & editing). A.V.V. (Data curation, Formal analysis, Investigation). L.C.K. (Data curation, Formal analysis, Investigation). G.K. (Investigation). A.S. (Investigation). A.S. (Investigation). T.S. (Data curation). A.I. (Validation). P.P.B. (Conceptualization, Data interpretation, Funding acquisition, Supervision, Validation, Writing—original draft, Writing—review & editing).

## Supplementary data

Supplementary Data are available at journal’s site Online.

## Data availability

Flow cytometry data will be uploaded to the FLOW repository once it becomes available again. Transcriptomic data used from (15) were deposited in Gene Expression Omnibus (GEO) under the accession code GSE145738. Custom code for transcriptomic analysis is available on GitHub (https://github.com/tschauer/Domino_RNAseq_2020). The mass spectrometry proteomics data have been deposited to the ProteomeXchange Consortium (http://proteomecentral.proteomexchange.org) via the PRIDE partner repository (46) with the dataset identifier PXD066107. Custom code for mass spectrometry analysis is available at https://github.com/anuroopv/RAmP. CUT&RUN sequencing data have been deposited in GEO under accession code GSE301786. Custom code for CUT&RUN and other analyses is available on Zenodo (https://doi.org/10.5281/zenodo.15844155).

## References

1. Chen, Y.C., Koutelou, E. and Dent, S.Y.R. (2022) Now open: Evolving insights to the roles of lysine acetylation in chromatin organization and function. Mol Cell, 82, 716–727.

2. Dhar, S., Gursoy-Yuzugullu, O., Parasuram, R. and Price, B.D. (2017) The tale of a tail: histone H4 acetylation and the repair of DNA breaks. Philos Trans R Soc Lond B Biol Sci, 372, 20160284.

3. Zaware, N. and Zhou, M.M. (2019) Bromodomain biology and drug discovery. Nat Struct Mol Biol, 26, 870–879.

4. Shvedunova, M. and Akhtar, A. (2022) Modulation of cellular processes by histone and non-histone protein acetylation. Nat Rev Mol Cell Biol, 23, 329–349.

5. Kamine, J., Elangovan, B., Subramanian, T., Coleman, D. and Chinnadurai, G. (1996) Identification of a cellular protein that specifically interacts with the essential cysteine region of the HIV-1 Tat transactivator. Virology, 216, 357–366.

6. Doyon, Y. and Cote, J. (2004) The highly conserved and multifunctional NuA4 HAT complex. Curr Opin Genet Dev, 14, 147–154.

7. Cheung, A.C.M. (2023) The NuA4 histone acetyltransferase: variations on a theme of SAGA. Nat Struct Mol Biol, 30, 1240–1241.

8. Sun, Y., Jiang, X. and Price, B.D. (2010) Tip60: connecting chromatin to DNA damage signaling. Cell Cycle, 9, 930–936.

9. Cheng, X., Cote, V. and Cote, J. (2021) NuA4 and SAGA acetyltransferase complexes cooperate for repair of DNA breaks by homologous recombination. PLoS Genet, 17, e1009459.

10. Giaimo, B.D., Ferrante, F., Herchenrother, A., Hake, S.B. and Borggrefe, T. (2019) The histone variant H2A.Z in gene regulation. Epigenetics Chromatin, 12, 37.

11. Altaf, M., Auger, A., Monnet-Saksouk, J., Brodeur, J., Piquet, S., Cramet, M., Bouchard, N., Lacoste, N., Utley, R.T., Gaudreau, L. et al. (2010) NuA4-dependent acetylation of nucleosomal histones H4 and H2A directly stimulates incorporation of H2A.Z by the SWR1 complex. J Biol Chem, 285, 15966–15977.

12. Cheng, X., Auger, A., Altaf, M., Drouin, S., Paquet, E., Utley, R.T., Robert, F. and Cote, J. (2015) Eaf1 Links the NuA4 Histone Acetyltransferase Complex to Htz1 Incorporation and Regulation of Purine Biosynthesis. Eukaryot Cell, 14, 535–544.

13. Auger, A., Galarneau, L., Altaf, M., Nourani, A., Doyon, Y., Utley, R.T., Cronier, D., Allard, S. and Cote, J. (2008) Eaf1 is the platform for NuA4 molecular assembly that evolutionarily links chromatin acetylation to ATP-dependent exchange of histone H2A variants. Mol Cell Biol, 28, 2257–2270.

14. Scacchetti, A. and Becker, P.B. (2021) Variation on a theme: Evolutionary strategies for H2A.Z exchange by SWR1-type remodelers. Curr Opin Cell Biol, 70, 1–9.

15. Scacchetti, A., Schauer, T., Reim, A., Apostolou, Z., Campos Sparr, A., Krause, S., Heun, P., Wierer, M. and Becker, P.B. (2020) Drosophila SWR1 and NuA4 complexes are defined by DOMINO isoforms. Elife, 9, e56325.

16. Ruhf, M.L., Braun, A., Papoulas, O., Tamkun, J.W., Randsholt, N. and Meister, M. (2001) The domino gene of Drosophila encodes novel members of the SWI2/SNF2 family of DNA-dependent ATPases, which contribute to the silencing of homeotic genes. Development, 128, 1429–1441.

17. Wichmann, J., Pitt, C., Eccles, S., Garnham, A.L., Li-Wai-Suen, C.S.N., May, R., Allan, E., Wilcox, S., Herold, M.J., Smyth, G.K. et al. (2022) Loss of TIP60 (KAT5) abolishes H2AZ lysine 7 acetylation and causes p53, INK4A, and ARF-independent cell cycle arrest. Cell Death Dis, 13, 627.

18. Martin, B.J.E., Ablondi, E.F., Goglia, C., Mimoso, C.A., Espinel-Cabrera, P.R. and Adelman, K. (2023) Global identification of SWI/SNF targets reveals compensation by EP400. Cell, 186, 5290–5307 e5226.

19. Fazzio, T.G., Huff, J.T. and Panning, B. (2008) Chromatin regulation Tip(60)s the balance in embryonic stem cell self-renewal. Cell Cycle, 7, 3302–3306.

20. Chan, H.M., Narita, M., Lowe, S.W. and Livingston, D.M. (2005) The p400 E1A-associated protein is a novel component of the p53--> p21 senescence pathway. Genes Dev, 19, 196–201.

21. Ravens, S., Yu, C., Ye, T., Stierle, M. and Tora, L. (2015) Tip60 complex binds to active Pol II promoters and a subset of enhancers and co-regulates the c-Myc network in mouse embryonic stem cells. Epigenetics Chromatin, 8, 45.

22. Sapountzi, V. and Cote, J. (2011) MYST-family histone acetyltransferases: beyond chromatin. Cell Mol Life Sci, 68, 1147–1156.

23. Borner, K. and Becker, P.B. (2016) Splice variants of the SWR1-type nucleosome remodeling factor Domino have distinct functions during Drosophila melanogaster oogenesis. Development, 143, 3154–3167.

24. Mendez, J. and Stillman, B. (2000) Chromatin association of human origin recognition complex, cdc6, and minichromosome maintenance proteins during the cell cycle: assembly of prereplication complexes in late mitosis. Mol Cell Biol, 20, 8602–8612.

25. Meers, M.P., Bryson, T.D., Henikoff, J.G. and Henikoff, S. (2019) Improved CUT&RUN chromatin profiling tools. Elife, 8, e46314.

26. Carlson, M. (2025) org.Dm.eg.db: Genome wide annotation for Fly.

27. Yu, G., Wang, L.G., Han, Y. and He, Q.Y. (2012) clusterProfiler: an R package for comparing biological themes among gene clusters. OMICS, 16, 284–287.

28. Dobin, A., Davis, C.A., Schlesinger, F., Drenkow, J., Zaleski, C., Jha, S., Batut, P., Chaisson, M. and Gingeras, T.R. (2013) STAR: ultrafast universal RNA-seq aligner. Bioinformatics, 29, 15–21.

29. Li, B. and Dewey, C.N. (2011) RSEM: accurate transcript quantification from RNA-Seq data with or without a reference genome. BMC Bioinformatics, 12, 323.

30. R_Core_Team. (2025) R: A Language and Environment for Statistical Computing. R Foundation for Statistical Computing, Vienna, Austria. URL https://www.R-project.org/.

31. Cox, J. and Mann, M. (2008) MaxQuant enables high peptide identification rates, individualized p.p.b.-range mass accuracies and proteome-wide protein quantification. Nat Biotechnol, 26, 1367–1372.

32. Ritchie, M.E., Phipson, B., Wu, D., Hu, Y., Law, C.W., Shi, W. and Smyth, G.K. (2015) limma powers differential expression analyses for RNA-sequencing and microarray studies. Nucleic Acids Res, 43, e47.

33. Hostrup, M., Lemminger, A.K., Stocks, B., Gonzalez-Franquesa, A., Larsen, J.K., Quesada, J.P., Thomassen, M., Weinert, B.T., Bangsbo, J. and Deshmukh, A.S. (2022) High-intensity interval training remodels the proteome and acetylome of human skeletal muscle. Elife, 11, e69802.

34. Xiao, Y., Hsiao, T.H., Suresh, U., Chen, H.I., Wu, X., Wolf, S.E. and Chen, Y. (2014) A novel significance score for gene selection and ranking. Bioinformatics, 30, 801–807.

35. Martin, M. Cutadapt removes adapter sequences from high-throughput sequencing reads. EMBnet.journal, 17, 10–12.

36. Langmead, B. and Salzberg, S.L. (2012) Fast gapped-read alignment with Bowtie 2. Nat Methods, 9, 357–359.

37. Langmead, B., Wilks, C., Antonescu, V. and Charles, R. (2019) Scaling read aligners to hundreds of threads on general-purpose processors. Bioinformatics, 35, 421–432.

38. Danecek, P., Bonfield, J.K., Liddle, J., Marshall, J., Ohan, V., Pollard, M.O., Whitwham, A., Keane, T., McCarthy, S.A., Davies, R.M. et al. (2021) Twelve years of SAMtools and BCFtools. Gigascience, 10, giab008.

39. Quinlan, A.R. and Hall, I.M. (2010) BEDTools: a flexible suite of utilities for comparing genomic features. Bioinformatics, 26, 841–842.

40. Ramirez, F., Ryan, D.P., Gruning, B., Bhardwaj, V., Kilpert, F., Richter, A.S., Heyne, S., Dundar, F. and Manke, T. (2016) deepTools2: a next generation web server for deep-sequencing data analysis. Nucleic Acids Res, 44, W160–165.

41. Zhang, Y., Liu, T., Meyer, C.A., Eeckhoute, J., Johnson, D.S., Bernstein, B.E., Nusbaum, C., Myers, R.M., Brown, M., Li, W. et al. (2008) Model-based analysis of ChIP-Seq (MACS). Genome Biol, 9, R137.

42. Kharchenko, P.V., Alekseyenko, A.A., Schwartz, Y.B., Minoda, A., Riddle, N.C., Ernst, J., Sabo, P.J., Larschan, E., Gorchakov, A.A., Gu, T. et al. (2011) Comprehensive analysis of the chromatin landscape in Drosophila melanogaster. Nature, 471, 480–485.

43. Ibarra-Morales, D., Rauer, M., Quarato, P., Rabbani, L., Zenk, F., Schulte-Sasse, M., Cardamone, F., Gomez-Auli, A., Cecere, G. and Iovino, N. (2021) Histone variant H2A.Z regulates zygotic genome activation. Nat Commun, 12, 7002.

44. Caron, C., Boyault, C. and Khochbin, S. (2005) Regulatory cross-talk between lysine acetylation and ubiquitination: role in the control of protein stability. Bioessays, 27, 408–415.

45. Wang, J., He, H., Chen, B., Jiang, G., Cao, L., Jiang, H., Zhang, G., Chen, J., Huang, J., Yang, B. et al. (2020) Acetylation of XPF by TIP60 facilitates XPF-ERCC1 complex assembly and activation. Nat Commun, 11, 786.

46. Ahmad, S., Cote, V., Cheng, X., Bourriquen, G., Sapountzi, V., Altaf, M. and Cote, J. (2021) Antagonistic relationship of NuA4 with the non-homologous end-joining machinery at DNA damage sites. PLoS Genet, 17, e1009816.

47. Wang, T., Zou, Y., Meng, H., Zheng, P., Teng, J., Huang, N. and Chen, J. (2024) Securin acetylation prevents precocious separase activation and premature sister chromatid separation. Curr Biol, 34, 1295–1308 e1295.

48. Sang, C., Li, X., Liu, J., Chen, Z., Xia, M., Yu, M. and Yu, W. (2024) Reversible acetylation of HDAC8 regulates cell cycle. EMBO Rep, 25, 3925–3943.

49. Jostes, S., Vardabasso, C., Dong, J., Carcamo, S., Singh, R., Phelps, R., Meadows, A., Grossi, E., Hasson, D. and Bernstein, E. (2024) H2A.Z chaperones converge on E2F target genes for melanoma cell proliferation. Genes Dev, 38, 336–353.

50. Kiss, A.E., Venkatasubramani, A.V., Pathirana, D., Krause, S., Sparr, A.C., Hasenauer, J., Imhof, A., Muller, M. and Becker, P.B. (2024) Processivity and specificity of histone acetylation by the male-specific lethal complex. Nucleic Acids Res, 52, 4889–4905.

51. Grienenberger, A., Miotto, B., Sagnier, T., Cavalli, G., Schramke, V., Geli, V., Mariol, M.C., Berenger, H., Graba, Y. and Pradel, J. (2002) The MYST domain acetyltransferase Chameau functions in epigenetic mechanisms of transcriptional repression. Curr Biol, 12, 762–766.

52. Peleg, S., Feller, C., Forne, I., Schiller, E., Sevin, D.C., Schauer, T., Regnard, C., Straub, T., Prestel, M., Klima, C. et al. (2016) Life span extension by targeting a link between metabolism and histone acetylation in Drosophila. EMBO Rep, 17, 455–469.

53. Venkatasubramani, A.V., Ichinose, T., Kanno, M., Forne, I., Tanimoto, H., Peleg, S. and Imhof, A. (2023) The fruit fly acetyltransferase chameau promotes starvation resilience at the expense of longevity. EMBO Rep, 24, e57023.

54. Yustis, J.C., Devoucoux, M. and Cote, J. (2024) The Functional Relationship Between RNA Splicing and the Chromatin Landscape. J Mol Biol, 436, 168614.

55. Bhatnagar, A., Krick, K., Karisetty, B.C., Armour, E.M., Heller, E.A. and Elefant, F. (2023) Tip60’s Novel RNA-Binding Function Modulates Alternative Splicing of Pre-mRNA Targets Implicated in Alzheimer’s Disease. J Neurosci, 43, 2398–2423.

56. Rust, K., Tiwari, M.D., Mishra, V.K., Grawe, F. and Wodarz, A. (2018) Myc and the Tip60 chromatin remodeling complex control neuroblast maintenance and polarity in Drosophila. EMBO J, 37, e98659.

57. Tue, N.T., Yoshioka, Y., Mizoguchi, M., Yoshida, H., Zurita, M. and Yamaguchi, M. (2017) DREF plays multiple roles during Drosophila development. Biochim Biophys Acta Gene Regul Mech, 1860, 705–712.

58. Hoareau, M., Rincheval-Arnold, A., Gaumer, S. and Guenal, I. (2024) DREAM a little dREAM of DRM: Model organisms and conservation of DREAM-like complexes: Model organisms uncover the mechanisms of DREAM-mediated transcription regulation. Bioessays, 46, e2300125.

59. Piunti, A. and Shilatifard, A. (2016) Epigenetic balance of gene expression by Polycomb and COMPASS families. Science, 352, aad9780.

60. Goodman, R.H. and Smolik, S. (2000) CBP/p300 in cell growth, transformation, and development. Genes Dev, 14, 1553–1577.

61. Huang, F., Abmayr, S.M. and Workman, J.L. (2016) Regulation of KAT6 Acetyltransferases and Their Roles in Cell Cycle Progression, Stem Cell Maintenance, and Human Disease. Mol Cell Biol, 36, 1900–1907.

62. Sheikh, B.N., Guhathakurta, S. and Akhtar, A. (2019) The non-specific lethal (NSL) complex at the crossroads of transcriptional control and cellular homeostasis. EMBO Rep, 20, e47630.

63. Golomb, L., Volarevic, S. and Oren, M. (2014) p53 and ribosome biogenesis stress: the essentials. FEBS Lett, 588, 2571–2579.

64. Huang, F., Saraf, A., Florens, L., Kusch, T., Swanson, S.K., Szerszen, L.T., Li, G., Dutta, A., Washburn, M.P., Abmayr, S.M. et al. (2016) The Enok acetyltransferase complex interacts with Elg1 and negatively regulates PCNA unloading to promote the G1/S transition. Genes Dev, 30, 1198–1210.

65. Flegel, K., Grushko, O., Bolin, K., Griggs, E. and Buttitta, L. (2016) Roles for the Histone Modifying and Exchange Complex NuA4 in Cell Cycle Progression in Drosophila melanogaster. Genetics, 203, 1265–1281.

66. Noguchi, C., Singh, T., Ziegler, M.A., Peake, J.D., Khair, L., Aza, A., Nakamura, T.M. and Noguchi, E. (2019) The NuA4 acetyltransferase and histone H4 acetylation promote replication recovery after topoisomerase I-poisoning. Epigenetics Chromatin, 12, 24.

67. Schleicher, E.M., Dhoonmoon, A., Jackson, L.M., Khatib, J.B., Nicolae, C.M. and Moldovan, G.L. (2022) The TIP60-ATM axis regulates replication fork stability in BRCA-deficient cells. Oncogenesis, 11, 33.

68. Duronio, R.J. and Marzluff, W.F. (2017) Coordinating cell cycle-regulated histone gene expression through assembly and function of the Histone Locus Body. RNA Biol, 14, 726–738.

69. Messina, G., Prozzillo, Y., Monache, F.D., Santopietro, M.V. and Dimitri, P. (2022) Evolutionary conserved relocation of chromatin remodeling complexes to the mitotic apparatus. BMC Biol, 20, 172.

