## Supplementary Figures for "The Tip60 acetylome is a hallmark of the proliferative state in *Drosophila*"

#### Supplementary Figure S1.

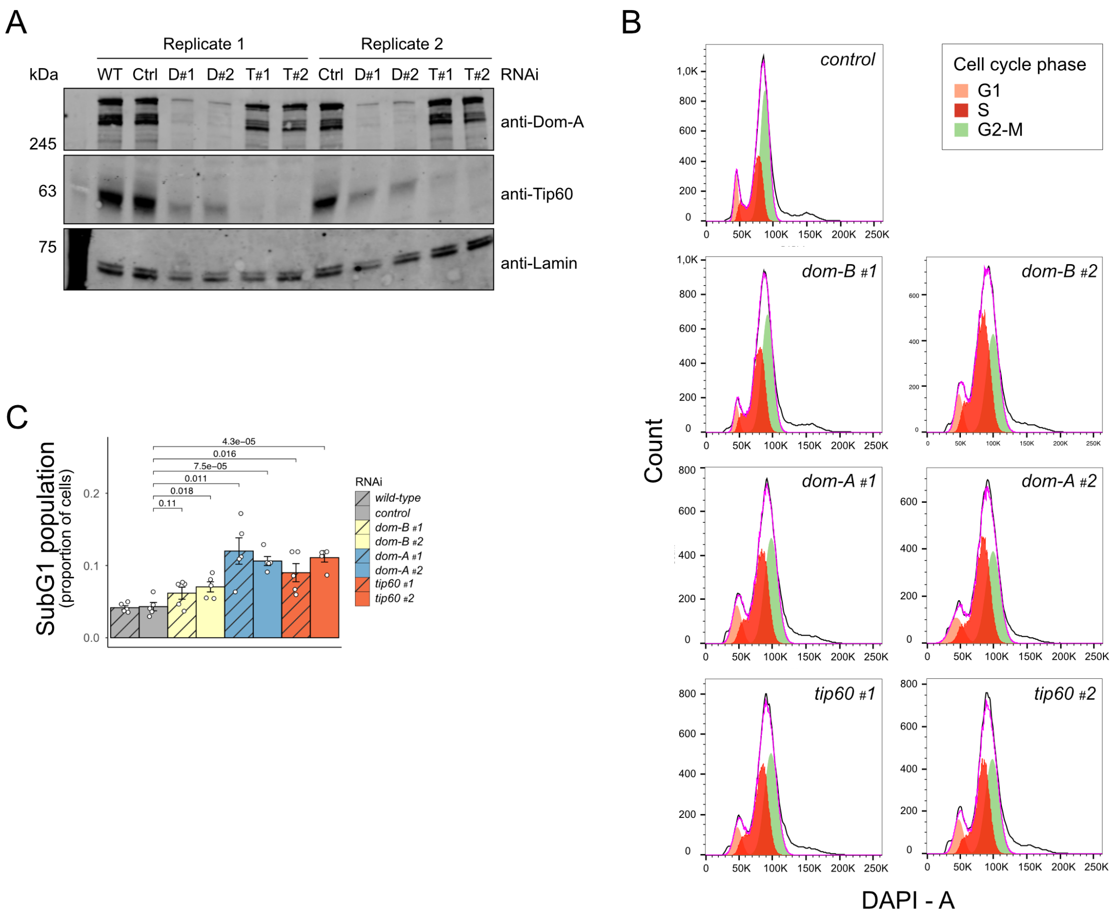


**Supplementary Figure S1. Depletion of DOM-A/ TIP60 leads to cell cycle arrest in Kc167 cells.**

A) Immunoblots of whole cell extracts showing the knockdown efficiency of *dom-A* and *tip60* in two different experimental replicates. Two different RNAi constructs were used throughout the analysis: D#1 and D#2 directed against *dom-A* mRNA and T#1 and T#2, directed against *tip60* mRNA. WT: naïve Kc167 cells. Ctrl: control RNAi directed against *gst*. Lamin serves as a loading control. The membranes were probed with antibodies indicated to the right. Note the reduction of Tip60 upon Dom-A-depletion, but not vice versa (1).

B) Cell cycle profiles of the same depleted cell populations shown in Figure 1D, with phase distributions determined using the Watson algorithm.

C) Quantification of the sub-G1 population from the cell cycle analysis in Figure 1D. The error bars indicate standard error of the mean (SEM) of 5 biological replicates.

#### Supplementary Figure S3.

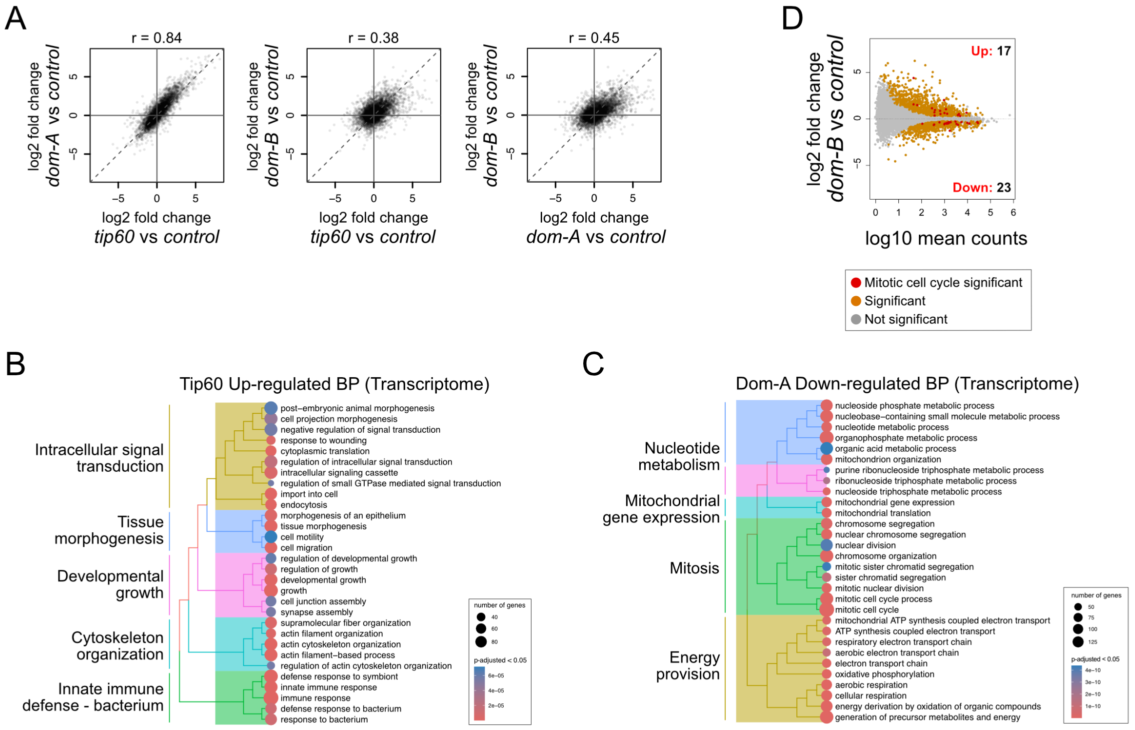


**Supplementary Figure S3: The DOM-A/TIP60 complex promotes transcription of cell cycle genes**

A) Scatter plots with pairwise comparisons of log2 fold gene expression changes in cells where Tip60, Dom-A, or Dom-B had been depleted. The Spearman correlation coefficient (r) is shown at the top of the plot.

B) Dendrogram depicting the top 30 significant GO terms from ORA of Tip60 upregulated biological processes of the transcriptome, filtered by p-adjusted cut-off < 0.05. Color of the circles indicates enrichment and the size indicates number of genes annotated with the corresponding pathway. GO terms were clustered based on semantic similarity and the terms that were represented the most within a cluster were mentioned.

C) Dendrogram depicting the top 30 significant GO terms from ORA of Dom-A downregulated biological processes of the transcriptome, filtered by p-adjusted cut-off < 0.05. Color of the circles indicates enrichment and the size indicates number of genes annotated with the corresponding pathway. GO terms were clustered based on semantic similarity and the terms that were represented the most within a cluster were mentioned.

D) MA plot comparing log2 fold-change of mean RNA-seq counts, derived from three independent experiments of control Kc167 cells and cells depleted of Dom-B. Differentially expressed genes within the indicated gene group are shown in red (p-adjusted < 0.05), differentially expressed genes but not within the corresponding gene group are shown in orange, and non-differentially expressed genes and not within the gene group are shown in grey.

Supplementary Figure S4.

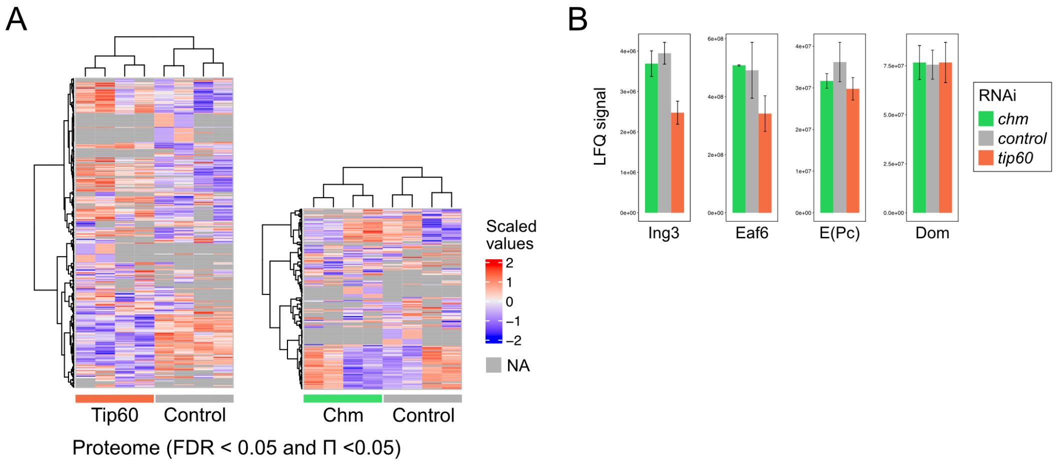


**Supplementary Figure S4. Deregulation of cell cycle proteins upon depletion of Tip60**

A) Heatmap displaying all significantly altered proteins (FDR < 0.05 and $\Pi < 0.05$) in Tip60-depleted versus control cells (left) and Chm-depleted versus control cells (right). Heatmap values represent scaled raw intensity measurements, with missing values (NAs) shown in gray.

D) LFQ signal intensities of selected subunits of the DOM-A/TIP60 complex upon depletion of Tip60 or Chm. The error bars indicate standard error (SE).

### Supplementary Figure S5.

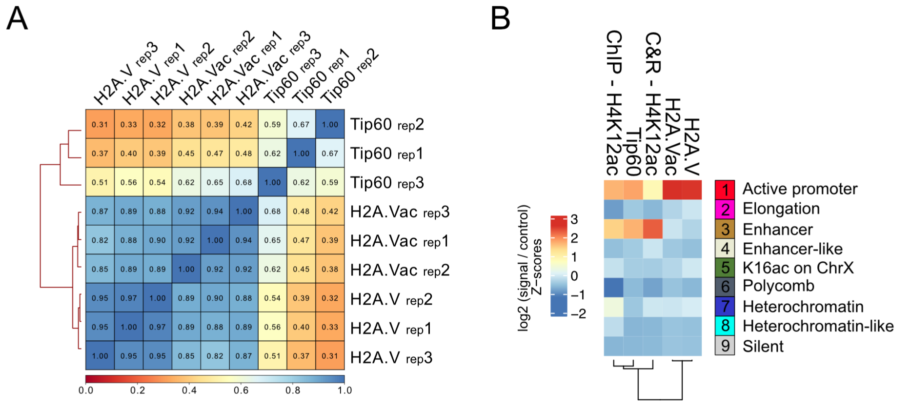


**Supplementary Figure S5. TIP60 binding at active promoters correlates with H2A.Vac.**

A) Hierarchical clustering of all CUT&RUN replicates for Tip60, H2A.V and acetylated H2A.V. The Spearman correlation coefficients were calculated based on read counts (scores) per 10 kb bin.

B) Chromatin state enrichment analysis as shown in Figure 4C with the addition of an H4K12ac ChIP-seq profile (n=1).

Supplementary Figure S6.

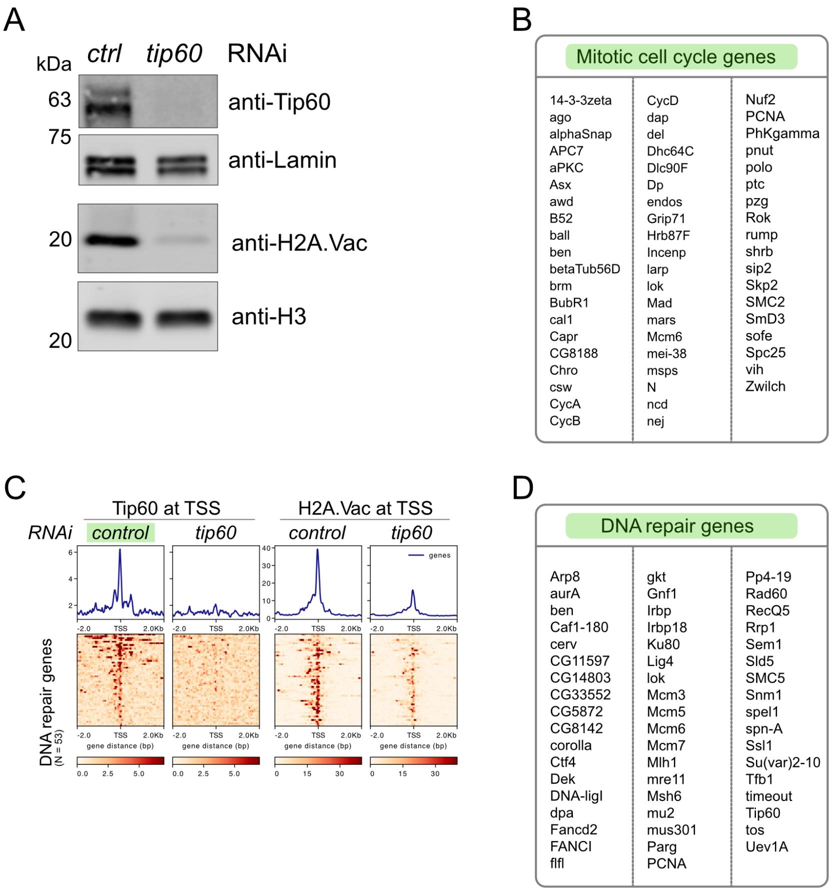


**Supplementary Figure S6. Depletion of TIP60 leads to reduced H2A.Vac.**

A) Immunoblotting of Tip60 and acetylated H2A.V in the chromatin fraction of control or *tip60* RNAi-treated Kc167 cells. Lamin and H3 serve as loading controls for Tip60 and acetylated H2A.V, respectively.

B) List of 'mitotic cell cycle genes' that show reduced expression in Tip60-depleted cells (p-adjusted < 0.05) and genomic binding on TSSs of Tip60.

C) Enrichment of Tip60 and H2A.Vac on genes of the GO-term "DNA repair" that show reduced expression in Tip60-depleted cells (p-adjusted < 0.05). Enrichment is illustrated by cumulative profiles (top) and heatmaps (bottom). Coverage windows of 4 kb around the TSS are selected and the mean is calculated for each column. Loci are sorted based on Tip60-control RNAi (colored in green). The coverage of each replicate was normalized to its corresponding IgG control (for H2A.V and H2A.Vac) or pre-immune serum (for Tip60), and the mean signal of 3 biological replicates was calculated.

D) List of DNA repair associated genes that show reduced expression in Tip60-depleted cells (p-adjusted < 0.05) and genomic binding on TSSs of Tip60.

Supplementary Figure S7.

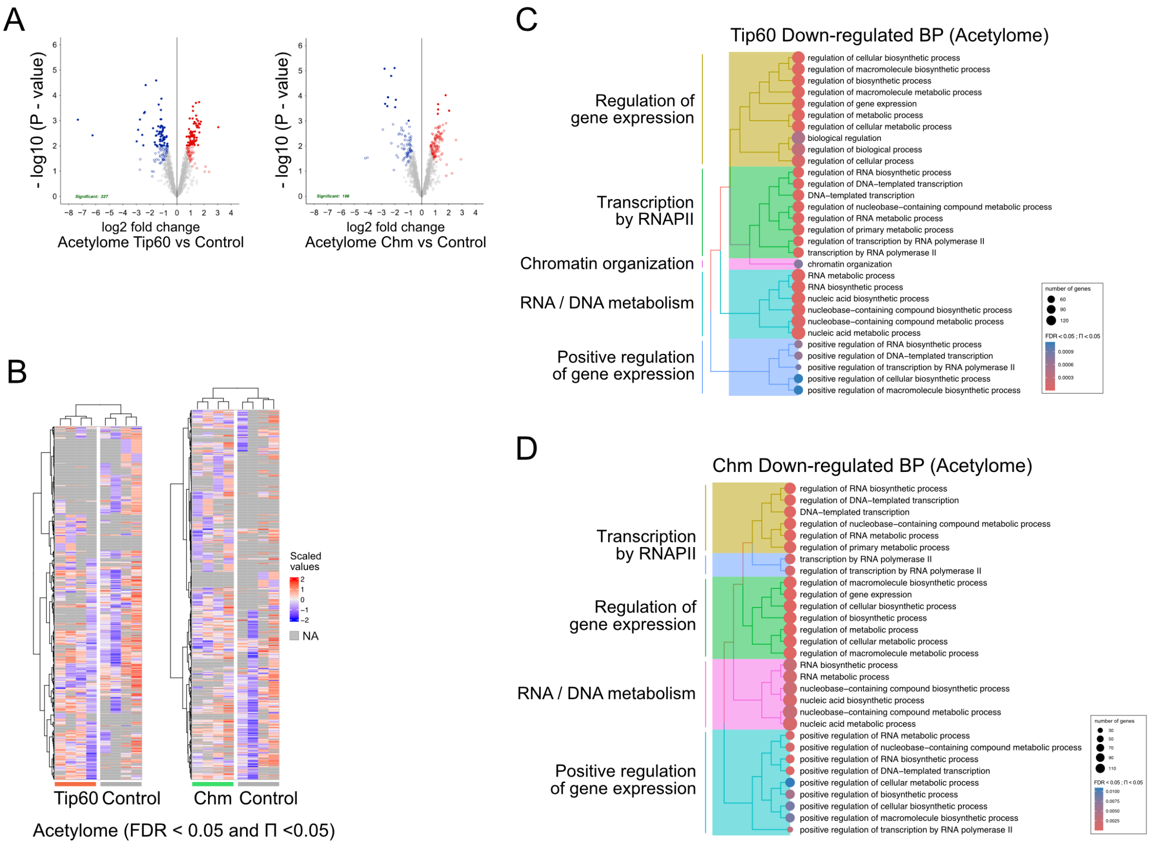


**Supplementary Figure S7. Depletion of TIP60 leads to numerous acetylome changes**

A) Volcano plots displaying the log2 fold-change (x-axis) versus the –log10 P-value (y-axis) for Class 2 acetylated peptides in Tip60-depleted versus control cells (left) and Chm-depleted versus control cells (right). Filled circles represent peptides with FDR < 0.05, while open circles indicate peptides with Π < 0.05. The color of each point reflects the direction and nature of differential regulation.

B) Heatmap depicting acetylated Class 1 peptides and significantly altered Class 2 peptides (FDR < 0.05 and Π < 0.05) in Tip60- and Chm-depleted cells compared to controls. Heatmap values correspond to scaled raw intensity measurements, with missing values (NA) shown in gray.

C) Dendrogram illustrating the top 30 significantly downregulated Gene Ontology (GO) Biological Process (BP) terms enriched among Class 1 and Class 2 acetylated proteins in the Tip60-depleted acetylome, based on over-representation analysis (ORA) filtered at FDR < 0.05 and Π < 0.05. Circle color denotes enrichment score, and size represents the number of genes associated with each GO term. GO terms were clustered by semantic similarity, with the most representative term for each cluster displayed.

D) Dendrogram equivalent to (C), summarizing the top 30 significantly downregulated GO BP terms in the Chm-depleted acetylome.

Reference

1. Scacchetti, A., Schauer, T., Reim, A., Apostolou, Z., Campos Sparr, A., Krause, S., Heun, P., Wierer, M. and Becker, P.B. (2020) Drosophila SWR1 and NuA4 complexes are defined by DOMINO isoforms. *Elife*, **9**, e56325.
